# LAP2alpha maintains a mobile and low assembly state of A-type lamins in the nuclear interior

**DOI:** 10.1101/2020.09.25.313296

**Authors:** Nana Naetar, Konstantina Georgiou, Christian Knapp, Irena Bronshtein, Elisabeth Zier, Petra Fichtinger, Thomas Dechat, Yuval Garini, Roland Foisner

## Abstract

Lamins form stable filaments at the nuclear periphery in metazoans. Unlike B-type lamins, lamins A and C localize also in the nuclear interior, where they interact with lamin-associated polypeptide 2 alpha (LAP2α). We show that lamin A in the nuclear interior is formed from newly expressed pre-lamin A during processing and from soluble mitotic mature lamins in a LAP2α-independent manner. Binding of LAP2α to lamins A/C in the nuclear interior during interphase inhibits formation of higher order structures of lamin A/C *in vitro* and *in vivo*, keeping lamin A/C in a mobile low assembly state independent of lamin A/C S22 phosphorylation. Loss of LAP2α causes formation of larger, less mobile and biochemically stable lamin A/C structures in the nuclear interior, which reduce the mobility of chromatin. We propose that LAP2α is essential to maintain a mobile lamin A/C pool in the nuclear interior, which is required for proper nuclear functions.

## Introduction

Lamins are intermediate filament proteins in metazoan nuclei that, together with numerous inner nuclear membrane proteins, form a filamentous protein meshwork at the nuclear periphery, called the nuclear lamina (Gruenbaum & Foisner, 2015). Based on their biochemical properties and expression patterns, lamins are grouped into two subtypes, A-type and B-type lamins. B-type lamins are ubiquitously expressed in all cell types and throughout development (Yang, Jung, Coffinier, Fong, & Young, 2011), whereas A-type lamins are expressed at low levels in embryonic stem cells and undifferentiated cells, but are significantly upregulated during differentiation (Constantinescu, Gray, Sammak, Schatten, & Csoka, 2006; Eckersley-Maslin, Bergmann, Lazar, & Spector, 2013; Rober, Weber, & Osborn, 1989). In mammalian cells, the major A-type lamins are lamins A and C encoded by the *Lmna* gene, whereas the two major B-type lamins, lamins B1 and B2, are encoded by *Lmnb1* and *Lmnb2*, respectively (Gruenbaum & Foisner, 2015). Both lamin subtypes share a similar intermediate filament protein-type domain structure with a central rod domain, an N-terminal head and a globular C-terminal tail, which contains a nuclear localization signal, an Ig fold and, except for lamin C, a C-terminal CaaX motif (C: cysteine; a: aliphatic amino acid; X: any amino acid) (de Leeuw, Gruenbaum, & Medalia, 2018; Gruenbaum & Medalia, 2015). The CaaX motif undergoes a series of post-translational modifications, including farnesylation of the cysteine, removal of the last three amino acids, followed by carboxymethylation (Rusinol & Sinensky, 2006). Whereas B-type lamins remain farnesylated and carboxymethylated, pre-lamin A undergoes an additional processing step catalyzed by the metalloprotease Zmpste24, leading to the removal of 15 amino acids from its C-terminus, including the farnesylated cysteine residue. As a consequence, mature B-type lamins are tightly associated with the nuclear membrane via their farnesylated C-terminus and mainly localize at the nuclear periphery, whereas mature lamin A is found both in filaments within the peripheral nuclear lamina, and additionally in a soluble and dynamic pool in the nuclear interior (Naetar, Ferraioli, & Foisner, 2017). Lamin C that lacks a CaaX motif contributes also to the peripheral and nucleoplasmic pool of A-type lamins.

The structure and functions of lamin filaments at the nuclear periphery are fairly well understood. Lamins form dimers via their central rod domains, which further assemble into head-to-tail polymers (de Leeuw et al., 2018). Recent work by the Medalia lab using cryo-electron tomography revealed that in mammalian nuclei two head-to-tail filaments assemble laterally into 3.5 nm thick filaments in a staggered fashion (Turgay et al., 2017). Lamin filaments at the nuclear periphery are considered stable, resistant to biochemical extraction and highly immobile (Bronshtein et al., 2015; Moir, Yoon, Khuon, & Goldman, 2000; Shimi et al., 2008). Together with proteins of the inner nuclear membrane, they fulfill essential functions, defining the mechanical properties of nuclei (Cho et al., 2019; Davidson & Lammerding, 2014; Osmanagic-Myers, Dechat, & Foisner, 2015) and regulating higher order chromatin organization through anchorage of peripheral heterochromatic genomic regions (van Steensel & Belmont, 2017).

The regulation and properties of A-type lamins in the nuclear interior are far less understood, although recent studies have unraveled novel functions of lamins in the nuclear interior in chromatin regulation that are fundamentally different from those fulfilled by the nuclear lamina (Gesson, Vidak, & Foisner, 2014; Naetar et al., 2017). Nucleoplasmic lamins A and C bind to and regulate euchromatic regions of the genome, globally affecting epigenetic modifications and possibly chromatin accessibility (Gesson et al., 2016; Naetar et al., 2017). They also provide chromatin “docking sites” in the nucleoplasm slowing down chromatin movement in nuclear space (Bronshtein et al., 2015). Moreover, intranuclear A-type lamins are required for the proper assembly of repressive polycomb protein foci (Bianchi et al., 2020; Cesarini et al., 2015) and are involved in the regulation of telomere function (Chojnowski et al., 2015; Gonzalez-Suarez et al., 2009; Wood et al., 2014). Nucleoplasmic lamins were also found to affect gene expression directly by binding to gene regulatory sequences, such as promoters and enhancers (Briand et al., 2018; Ikegami, Secchia, Almakki, Lieb, & Moskowitz, 2020; Oldenburg et al., 2017).

The structure and assembly state of nucleoplasmic lamins remain enigmatic. Fluorescence recovery after photobleaching (FRAP), continuous photobleaching (CP) and fluorescence correlation spectroscopy (FCS) studies of fluorescently tagged A-type lamins showed that the majority of nucleoplasmic lamin complexes are highly mobile compared to the stable peripheral lamina (Bronshtein et al., 2015; Moir et al., 2000; Shimi et al., 2008). Biochemical studies revealed that intranuclear lamins can be easily extracted in salt and detergent-containing buffers, suggesting that they have a low assembly state likely comprising dimers and/or short polymers (Kolb, Maass, Hergt, Aebi, & Herrmann, 2011; Naetar et al., 2008). Taken together, the dynamic nucleoplasmic lamin pool has very distinct properties compared to the static and stable peripheral lamina, allowing them to fulfill a unique set of functions. However, why and how nucleoplasmic A-type lamins display such different properties in the nuclear interior compared to their counterparts at the nuclear lamina remains poorly understood. Possible mechanisms include post-translational modifications and/or interactions with specific nucleoplasmic binding partners. For example, phosphorylation of lamins influences the localization, solubility and mobility of lamins both during mitosis and interphase (Kochin et al., 2014; Machowska, Piekarowicz, & Rzepecki, 2015). Moreover, the specific interaction partner of nucleoplasmic lamins, lamin-associated polypeptide 2 alpha (LAP2α) was suggested to regulate the localization of lamin A/C in the nuclear interior, since the nucleoplasmic lamin pool was found significantly reduced in the absence of LAP2α (Dechat et al., 2000; Gesson et al., 2016; Gesson et al., 2014; Naetar et al., 2008). However, the mechanisms remain unknown.

Here we address the open question if and how LAP2α aids nucleoplasmic A-type lamins to maintain their unique dynamic properties. We show that, unlike previously thought, the absence of LAP2α does not eliminate the nucleoplasmic lamin pool. Instead, nucloplasmic lamins become significantly more stable in the absence of LAP2α, possibly forming higher order structures in the nucleoplasm. The interaction of soluble lamins with LAP2α maintains their mobile, low assembly state in the nuclear interior, likely by directly inhibiting lamin assembly. This has direct functional consequences for the lamin-mediated regulation of chromatin mobility, slowing down chromatin diffusion in the absence of LAP2α.

## Results

### The nucleoplasmic lamin pool forms independently of LAP2α through at least two different processes

In order to elucidate the source of the nucleoplasmic pool of A-type lamins and the potential role of LAP2α in its formation we performed live cell imaging of wildtype and LAP2α knockout HeLa cells (generated by CRISPR-Cas9, see Fig. S1). We expressed GFP-pre-lamin A and imaged emerging pre-lamin A structures 5 hours post transfection, when the newly expressed pre-lamin A undergoes post-translational processing (Fig. 1A, see also video files 1 and 2 associated with Fig.1). The nascent, transiently farnesylated pre-lamin A initially localizes to the nuclear periphery with very little visible nucleoplasmic staining (Fig. 1A, 0-20’). Time-resolved quantification of the nucleoplasmic over peripheral fluorescence signal (Fig. 1A, lower panel) reveals that lamins in the nuclear interior emerge gradually within 20 minutes, probably after their release from the nuclear periphery during further processing and removal of the C-terminal farnesyl group. Accordingly, an ectopically expressed, farnesylation-deficient mature lamin A (lacking its C-terminal 15 amino acids) is detectable prominently in the nucleoplasm at early timepoints when lamin structures first emerge (Fig. S2A, compare columns 1 and 2, see also video file 5 associated with Fig. S2). Furthermore, an assembly-deficient lamin AΔK32 mutant (Bank et al., 2011; Bertrand et al., 2012) expressed in its pre-lamin form appears first at the nuclear periphery, before it translocates completely into the nucleoplasm, while the mature form of lamin AΔK32 mutant is exclusively in the nuclear interior throughout imaging (Fig. S2A, compare columns 3 and 4, see also video files 6 and 7 associated with Fig. S2). These data suggest that post-translational processing rather than filament assembly is the driving force for the initial localization of nascent pre-lamin A to the nuclear periphery.

**Figure 1.**
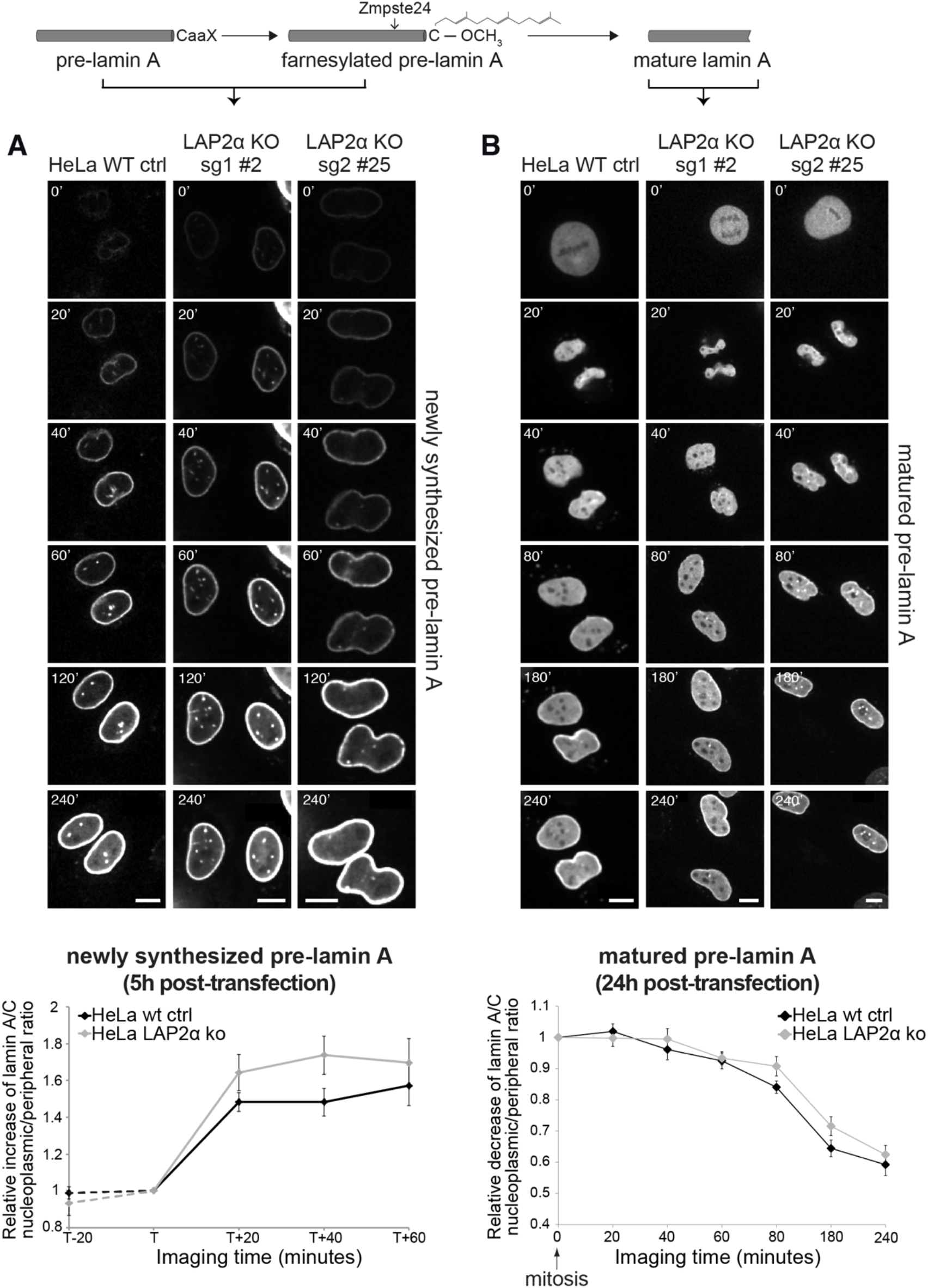
Absence of LAP2α does not affect formation of a nucleoplasmic pool of exogenously expressed GFP-lamin. **A.** Wildtype (WT) and LAP2α knockout (KO) HeLa cells (see Fig. S1) were transiently transfected with EGFP-pre-lamin A and analyzed by live-cell imaging either 5 hours **(A)** or 24 hours **(B)** post-transfection. Schematic drawing on top explains lamin A processing state at the time of imaging. Time is indicated in minutes in (A) and (B). Scale bar: 10 μm. See also video files: Videos 1 and 2, corresponding to panels 1 and 2 in (A) and videos 3 and 4, corresponding to panels 1 and 2 in (B). Bottom: Ratio of nucleoplasmic to peripheral mean signal intensity of EGFP-lamin A was calculated and normalized to time point “T” in (A). “T” defines the moment preceding a significant increase of nucleoplasmic lamin A in each cell. “T+/−x” indicates imaging time relative to “T” with ‘x’ corresponding to the number of minutes. Dashed lines indicate measurements performed in a smaller number of cells (n_wt_=6, n_ko_=6), demonstrating that nucleoplasmic lamin levels remain steadily low prior to the initial release of lamin A into the nucleoplasm (time point 0’). In (B) the nucleoplasmic over peripheral signal ratio was normalized to mitosis/early G1 (time point 0’) and plotted versus imaging time. Graphs display mean values ± S.E.M. n_wt_=18, n_ko_=23 for (A) and n_wt_=10, n_ko_=12 for (B). p values (repeated measures 2-way ANOVA) are p_time_<0.0001 and p_genotype_= 0.2160 (non-significant) for (A) and p_time_<0.0001 and p_genotype_= 0.2564 (non-significant) for (B).

A fraction of the peripheral lamin A may then move into the nuclear interior by unknown mechanisms. In order to test whether LAP2α is involved in this process, we imaged nascent GFP-pre-lamin A in LAP2α knockout cells and determined the nucleoplasmic over peripheral fluorescence signal over time after first appearance of nascent pre-lamin A at the nuclear periphery (Fig. 1A). The increase in the lamin A signal in the nuclear interior occurred with similar kinetics in LAP2α knockout and wildtype cells.

To follow the dynamic behavior of fully processed, mature lamin A during the cell cycle, we expressed GFP-pre-lamin A in wildtype and LAP2α knockout HeLa cells and analyzed cells 24 hours post transfection, when the majority of the ectopic pre-lamin A has been processed into the mature form (Fig. 1B, see also video files 3 and 4 associated with Fig. 1). As previously reported (Broers et al., 1999; Moir et al., 2000), lamin A structures completely disassemble at the onset of mitosis (Fig. 1B, 0’). During nuclear reassembly, lamin A disperses uniformly throughout the nucleus in late telophase / early G1 phase (Fig. 1B, 20’). Lamin A levels in the nuclear interior then gradually decrease within around 200 minutes due to relocation of the majority of lamins to the nuclear periphery, but a fraction of lamin A remains in the nuclear interior throughout interphase (Fig.1B, 40-240’, see also (Dechat et al., 2000; Naetar et al., 2008). As expected, ectopically expressed mature GFP-lamin A shows the same cell cycle-dependent behavior (Fig. S2B, see also video files 8-10 associated with Fig. S2). However, an assembly-deficient mutant, lamin AΔK32 remains fully nucleoplasmic throughout G1 phase, independent of whether the pre- or mature form was expressed, Thus, lamin filament assembly is essential for the accumulation of A-type lamins at the nuclear periphery during G1 phase. Next, we performed live cell imaging of LAP2α knockout cells, to test whether LAP2α affects the redistribution of lamins from the nuclear interior to the periphery. Quantification of the nucleoplasmic over peripheral GFP-lamin A signal after mitosis revealed a similar gradual decrease in nucleoplasmic staining during G1 phase in LAP2α knockout and wildtype cells, and more surprisingly, the steady state levels of lamin A in the nuclear interior throughout interphase were unaffected in LAP2α knockout versus wildtype cells (Fig. 1B, lower panel).

In conclusion, we identified two sources of nucleoplasmic A-type lamins, i) newly synthesized lamin A that redistributes from the periphery to the nuclear interior upon post-translational processing, and ii) soluble A-type lamins of the previous mitosis that remain in the nucleoplasm during lamina assembly in G1 phase. However, none of these processes were significantly altered in the absence of LAP2α, suggesting that nucleoplasmic A-type lamins can be generated independently of LAP2α. In addition, steady-state lamin A levels in the nuclear interior in interphase apparently were unchanged in the absence of LAP2α, contrary to previous reports (Naetar et al., 2008).

### LAP2α-deficiency does neither affect the formation nor the steady state level of endogenous lamin A in the nuclear interior

We reasoned that the lack of any effect of LAP2α loss on the formation of the nucleoplasmic lamin A pool may be due to overexpression of the ectopic GFP-lamin A. Thus, in order to create a more physiological setting, we tagged the endogenous lamin A gene with mEos3.2, a monomeric variant of the photoconverter EosFP (Zhang et al., 2012), in wildtype and LAP2α knockout mouse fibroblasts using CRISPR-Cas9 (Fig. 2A, Fig. S3 and S4). Following a rigid testing of the clones (see details in Fig. S3), we picked a wildtype clone (WT clone #21) for further analyses, which displays normal nuclear morphology and harbors both, tagged and untagged lamin A/C (Fig. 2B, C and Fig. S3F, G). In addition, we generated isogenic LAP2α knockout clones from this WT clone by CRISPR-Cas9 (for details see Fig. S4). mEos3.2-tagged lamin A/C displayed normal localization at the nuclear periphery and in the nuclear interior in LAP2α wildtype and knockout cells (Fig. 2C) and was stably integrated into the nuclear lamina, as demonstrated by photoconversion of tagged lamins, where no significant mobility of photo-converted lamins in the lamina was observed for several hours (Fig. S3H).

**Figure 2.**
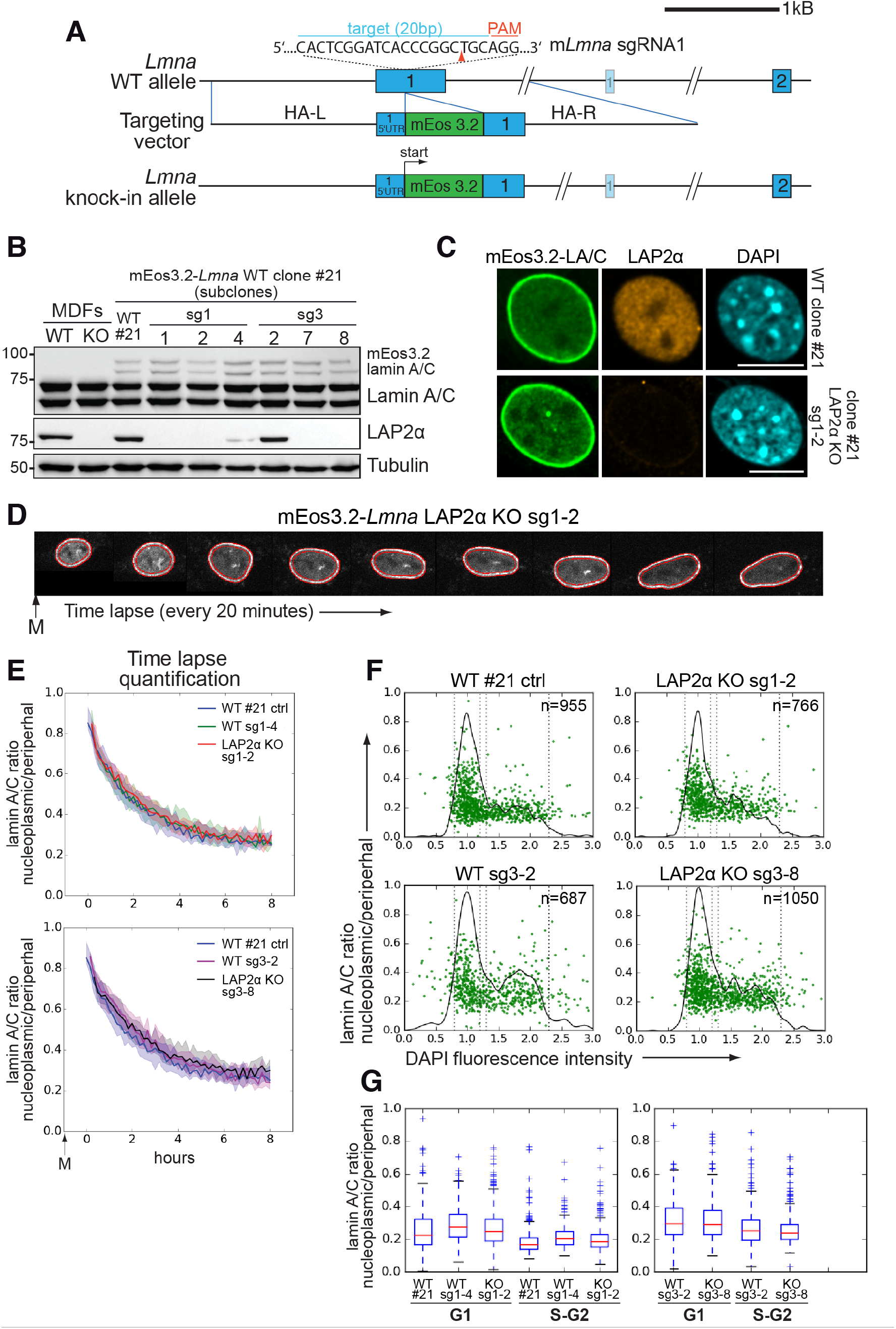
Absence of LAP2α does not alter endogenous nucleoplasmic lamin A/C levels. **(A)** Schematic view of exons 1 and 2 (bars) and adjacent introns of the mouse *Lmna* locus (top), the targeting construct provided as homology-directed repair template after Cas9-mediated double strand break (middle), and the knock-in allele after successful integration of mEos3.2 (bottom). Very long introns are not displayed in their original length as indicated by a double slash. The second, light-colored exon 1 encodes the N-terminus of meiosis-specific lamin C2. The target sequence of *Lmna*-specific sgRNA1 in exon 1 is shown (blue). Protospacer-adjacent motif (PAM) is marked in red. Red arrowhead: expected Cas9 cut site. HA-L: Homology Arm-Left; HA-R: Homology Arm-Right. For further cell characterization see Fig. S3. **(B)** mEos3.2-*Lmna* clone #21 derived from wildtype (WT) mouse dermal fibroblasts (MDFs) and subclones generated after treatment with LAP2α-specific sgRNAs 1 or 3 (see Fig. S4) were processed for Western blotting using antibodies to the indicated antigens (anti lamin A/C E1, anti LAP2α 1H11). Positions of mEos-tagged and untagged lamins A and C are indicated. Wildtype and LAP2α knockout (KO) MDFs were added as controls. **(C)** mEos3.2-*Lmna* WT clone #21 and LAP2α KO sg1-2 subclone were processed for immunofluorescence microscopy using antibodies against LAP2α (1H11) and DAPI to visualize DNA. Confocal images are shown. Bar: 10 μm. **(D)** Representative example of a cell expressing mEos3.2-lamin A/C tracked by live-cell imaging throughout G1, followed by automated calculation of nucleoplasmic over peripheral lamin A/C ratio (t_0_: first available image after mitosis). The area corresponding to the nuclear lamina is delineated by red lines and was defined by a custom-made algorithm implemented in FIJI/ImageJ software. Average values for isogenic mEos3.2-*Lmna* LAP2α WT and KO clones were plotted in curves **(E)**. The 20^th^ and 80^th^ percentile are displayed for each curve (shaded area). n_WT#21_=129, n_sg1-4_=159, n_sg1-2_=106, n_sg3-2_=111, n_sg3-8_=147 **(F)** Isogenic clones were fixed and DNA was stained with DAPI. Nucleoplasmic (N) to peripheral (P) signal intensity of mEos3.2-Lamin A/C was calculated for single cells (green dots) using a custom-made FIJI plugin and plotted versus DAPI fluorescence intensity, indicative of cell cycle stage. Black line outlines number of cells versus DAPI intensity (histogram). Vertical dotted lines indicate what was classified as G1 and S-G2 in (G). **(G)** Single cell N/P ratios from (E) were averaged for G1 and S/G2 and are shown in a box plot (median in red within the first and third quartiles; whiskers: minimal and maximal datapoint excluding outliers).

To investigate the potential role of LAP2α in the retention of endogenous lamins A and C in the nucleoplasm in G1 phase, or in the regulation of steady-state levels of lamin A/C in the nuclear interior throughout interphase, we followed mEos3.2-tagged lamin A/C in wildtype and LAP2α knockout isogenic clones by live cell imaging. We performed time lapse studies starting at the exit from mitosis through G1 phase, and determined the nucleoplasmic to peripheral lamin A/C signal ratio in hundreds of cells by automated quantification using a FIJI plugin (Figure 2D). Neither the post-mitotic dynamics of nucleoplasmic lamin localization nor its steady-state levels in late G1 phase were affected in LAP2α knockout versus wildtype cells (Fig. 2E).

To further analyze nucleoplasmic lamin A/C levels in interphase, we fixed mEos3.2-lamin A/C expressing wildtype and LAP2α knockout cells and automatically quantified their nucleoplasmic to peripheral lamin A/C ratio as described before. The ratio was then plotted against the intensity of the DAPI signal to stage cells within the cell cycle (Roukos, Pegoraro, Voss, & Misteli, 2015) (Fig. 2F). As expected, the nucleoplasmic to peripheral lamin A/C ratio was higher in G1 phase than in S phase, but both of these ratios were unchanged in the absence of LAP2α when compared to wildtype cells (Fig. 2F and G).

Thus, endogenous lamins A and C are retained in the nucleoplasm after mitosis and remain in the nuclear interior throughout interphase independent of LAP2α, which is in stark contrast to previous findings demonstrating a clear reduction of nucleoplasmic lamin A/C signal in LAP2α knockout cells and tissues by immunofluorescence microscopy (Naetar et al., 2008).

### LAP2α loss changes properties of nucleoplasmic lamin A/C rendering them less mobile and more resistant to biochemical extraction

In order to resolve these contradicting results, we first tested mEos3.2-lamin A/C expressing wildtype cells (WT#21) and isogenic LAP2α knockout clones by immunofluorescence microscopy using an antibody to the lamin A/C N-terminus, which, unlike an antibody to the lamin A/C C-terminus (see Fig S5C), preferentially stains nucleoplasmic lamins A and C (Gesson et al., 2016) (Fig. 3A and B, Fig. S5A). While the intranuclear mEos3.2 fluorescence signal was similar in LAP2α knockout and wildtype cells, the nucleoplasmic lamin antibody-derived signal was clearly reduced in LAP2α knockout versus wildtype cells (Fig. 3A, B and Fig. S5A, compare red and green channels and see quantification in Fig. 3D). Interestingly, the intranuclear antibody staining in LAP2α knockout cells was completely recovered following post-fixation treatment with DNase and RNase (Fig. 3C and D), suggesting epitope masking specifically in LAP2α knockout cells. In order to exclude the possibility that mEos3.2-lamin A/C protein behaves differently as compared to untagged lamin A/C (both are present in the WT#21 cell clones), we also tested cell clones expressing exclusively mEos3.2-tagged lamin A/C, without any untagged lamin A/C (see Fig. S5B, WT clone #5 and LAP2α KO clone #17). These clones displayed the same reduction in antibody-derived nucleoplasmic lamin staining in the absence of LAP2α, while the mEos3.2-lamin A/C fluorescence signal in the nuclear interior was unchanged (Fig. S5B).

**Figure 3.**
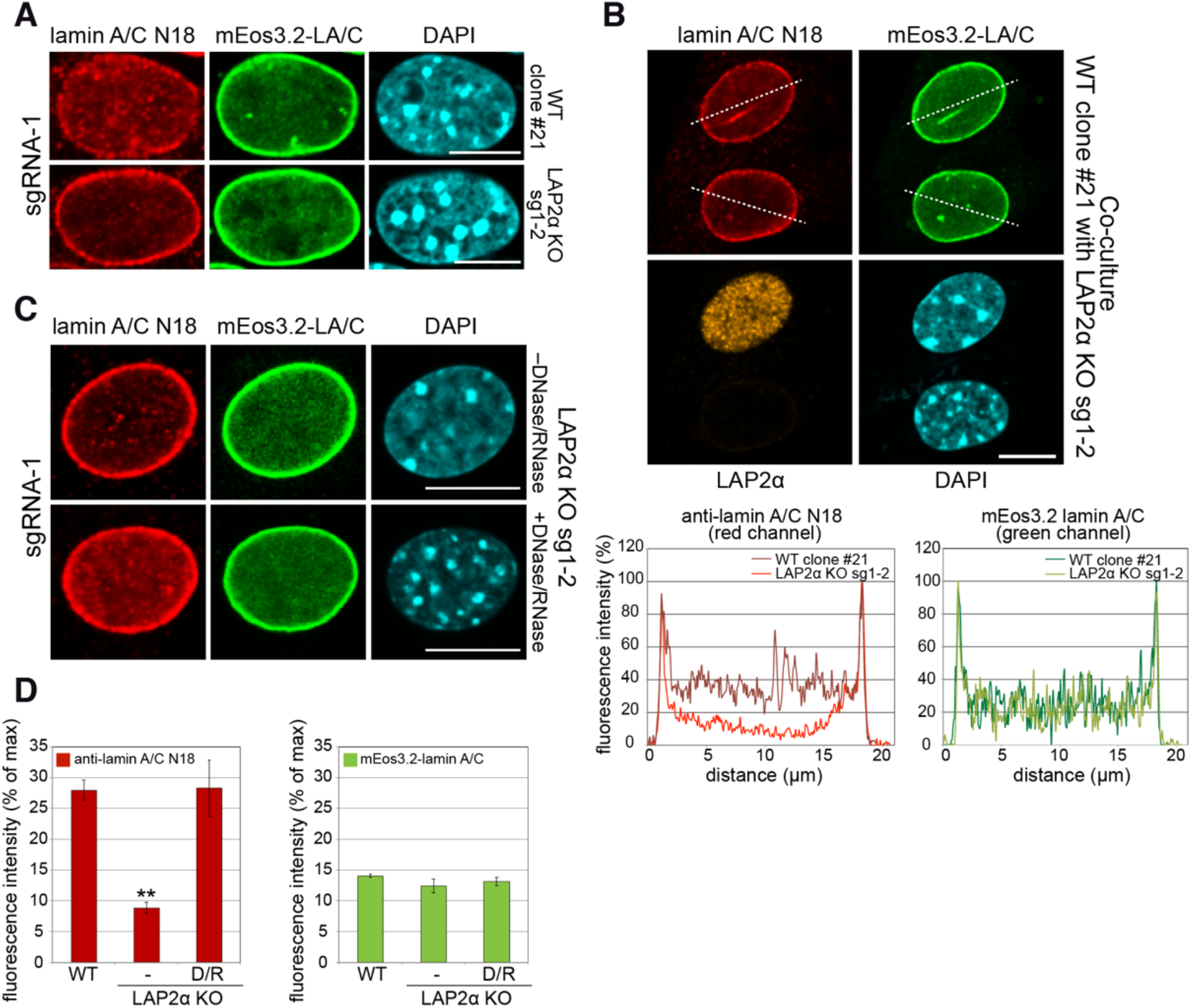
Absence of LAP2α reduces nucleoplasmic lamin A/C staining in immunofluorescence. **(A)** mEos3.2-*Lmna* WT clone #21 and LAP2α KO sg1-2 cells were processed for immunofluorescence microscopy using antibody N18 against lamin A/C, and DAPI to visualize DNA. Lamin A/C antibody N18 preferably stains nucleoplasmic lamins A/C. **(B)** mEos3.2-*Lmna* WT clone #21 and LAP2α KO sg1-2 cells were co-cultured and processed for immunofluorescence microscopy as in (A) using antibodies N18 against lamin A/C and 1H11 against LAP2α, and DAPI to visualize DNA. Fluorescence intensity was determined for each cell along the dashed line in the red and green channel and is depicted in percent of maximum value in the graphs below. **(C)** LAP2α KO sg1-2 cells were processed as in (A) using lamin A/C antibody N18 without or with prior treatment with DNase I and RNase A to reverse N18 epitope masking. Bar: 10 μm. **(D)** Average nucleoplasmic fluorescence intensity from 3 wildtype and 3 LAP2α KO clones with and without prior DNase/RNase (D/R) treatment for N18 stainings and for mEos3.2 lamin A/C signal is depicted. Data represent averages ± S.E.M. One-way ANOVA p value for N18 is 0.0020, F= 20.85; **p<0.01 (Tukey’s post-hoc test: WT vs. KO: p = 0.0034; KO vs KO plus DNase/RNase: p = 0.0033).

Overall, these data suggest that – instead of a complete absence of nucleoplasmic A-type lamins – the reduced staining in LAP2α knockout cells reflects an alteration of nucleoplasmic lamin A/C properties that led to masking of the N-terminal epitope recognized by the antibody.

To test potential changes in nucleoplasmic lamin A/C properties in LAP2α knockout versus wildtype cells, we first tested lamin mobility. While lamins at the periphery form stable filaments, nucleoplasmic lamins A/C were shown to be highly mobile (Broers et al., 1999; Kolb et al., 2011; Shimi et al., 2008). We tested the mobility of nucleoplasmic lamins by Continuous Photobleaching (CP), where the fluorescence intensity is measured in a small spot within the nucleoplasm of mEos3.2-lamin A/C expressing cells at a high frequency over time. It allows to reveal two fluorescent sub-populations with different mobility: an immobile fast bleaching fraction, and a highly mobile slow bleaching fraction (Fig. 4A, left panel). As previously demonstrated (Bronshtein et al., 2015), 10-40% of lamin A in the nucleoplasm is immobile in the majority of wildtype cells (Fig. 4A). Intriguingly, in the absence of LAP2α we observed a noticeable shift towards a higher fraction of immobile lamin A/C in the nuclear interior in the range of 30 up to 60% (Fig. 4A, right panel). Thus, the mobile fraction of lamin A in the nuclear interior is reduced in LAP2α knockout versus wildtype cells. In addition, we measured the mobile fraction by fluorescence correlation spectroscopy (FCS), which has previously revealed a slower and a faster-moving fraction within the mobile lamin A pool in the nuclear interior (Shimi et al., 2008). Despite the overall decrease in mobile lamin A/C in LAP2α knockout cells, we observed an increase in the diffusion coefficient of the remaining slow and fast-moving mobile lamin A/C fraction (Shimi et al., 2008), likely reflecting loss of binding to LAP2α (Fig. 4B).

**Figure 4.**
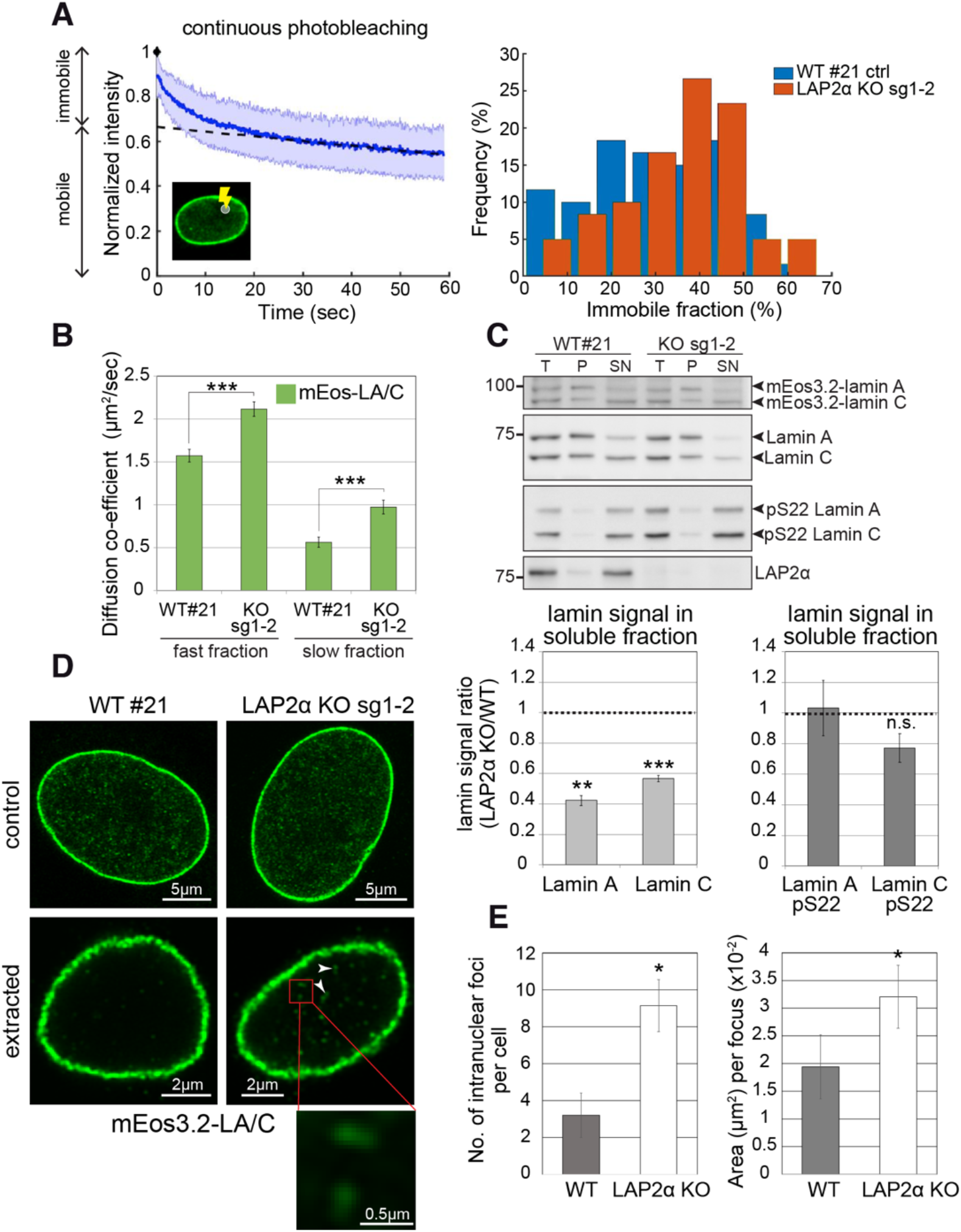
Nucleoplasmic lamin A/C are less mobile, more extraction-resistant and form larger assemblies in the absence of LAP2α. **(A)** Left: Depicted is a representative curve obtained by continuous photobleaching (CP, see schematic inset on the lower left) using WT clone #21 cells, allowing to determine mobile and immobile fractions (shown on the left side of the graph) of the measured mEos3.2-lamin A/C. Blue line represents normalized average intensity of mEos3.2-lamin A/C measured over time from a selected spot inside the nucleus. Right: Immobile fractions of mEos3.2-lamin A and C protein as calculated from measured CP curves of WT clone #21 and LAP2α KO sg1-2 cells are depicted as histogram. n_WT#21_=59, n_sg1-2_=60, p = 0.0053 (two-tailed student’s t test of arcsin transformed values) **(B)** Graph displays the diffusion co-efficient of fast and slow moving mEos3.2-lamins A and C as determined by fluorescence correlation spectroscopy (FCS) measurements. Data represent averages ± S.E.M.; n_WT#21 (fast fraction)_=106, n_WT#21 (slow fraction)_=96, n_sg1-2 (fast fraction)_=97, n_sg1-2 (slow fraction)_=99; ***p < 0.0001 (Mann-Whitney U test). **(C)** Cells from isogenic WT and LAP2α KO clones were extracted in salt and detergent-containing buffer (500 mM NaCl, 0.5% NP-40). Extracts were processed for Western blot analysis. Upper panel shows representative Western blots from WT#21 and LAP2α KO sg1-2 cells using antibodies against total lamin A/C (E1), lamin A/C phosphorylated at serine 22 (pS22) or LAP2α (1H11). T: total lysate; P: insoluble pellet fraction; SN: soluble, extracted supernatant fraction. Western blots were quantified and lamin signal in the supernatant was normalized to total lamin A/C. Graphs display lamins A and C levels in the supernatant fraction of LAP2α knockout samples as average fold difference ± S.E.M over wildtype samples. n_WT_=6, n_ko_=6; **p=0.00013, ***p=2.09E-5, n.s.: non-significant (p=0.072) (paired student’s t-test on log transformed values). **(D)** WT clone #21 and LAP2α KO sg1-2 cells were processed for immunofluorescence microscopy with and without prior extraction in salt and detergent-containing buffer. Confocal super-resolution (Airyscan) images of mEos3.2-lamin A/C signal are depicted. Larger lamin A/C nucleoplasmic assemblies in LAP2α knockout cells are marked by arrowheads and displayed as larger inset (bottom). **(E)** Graphs show quantification of number (No.) of intranuclear lamin A/C structures per cell and mean area per structure. Data represent averages ± S.E.M. Left graph: n_WT_ =5, n_KO_=7, p=0.0125 (unpaired two-tailed student’s t test); right graph: n_WT_ =16, n_KO_=64, p=0.0433 (Mann-Whitney U test); *p < 0.05.

To reveal further changes in lamin A properties in LAP2α knockout versus wildtype cells we performed biochemical extraction in buffers containing detergent and 150-500 mM salt. Lamins A/C were significantly less extractable in the absence of LAP2α in both, mouse fibroblasts and HeLa cells (Fig 4C and Fig. S6A), consistent with a shift of nucleoplasmic lamins into more stable, possibly higher assembly structures upon loss of LAP2α. Next, we aimed at visualizing the potentially higher assembly structures in LAP2α knockout cells by high resolution microscopy. While microscopy of untreated cells did not reveal clear differences in the appearance of nucleoplasmic lamin structures (Fig. 4D, upper panel), extraction of cells in detergent and salt-containing buffers prior to fixation revealed significantly more and larger, extraction resistant lamin A structures in the nuclear interior in LAP2α knockout versus wildtype mEos3.2-lamin A/C cells (Fig. 4D and quantification in Fig. 4E). Strikingly, while in wildtype cells these structures appeared as few punctae, LAP2α knockout cells displayed significantly more stable structures of an extended, possibly filamentous appearance (Figs. 4D, 4E). Altogether, loss of LAP2α does not lead to a reduction of the nucleoplasmic pool of lamin A/C, it rather affects its mobility and assembly state.

### LAP2α binds to intranuclear lamin A/C and inhibits its assembly

In order to better understand how LAP2α might influence the mobility and/or assembly state of nucleoplasmic lamin A/C, we performed *in vitro* interaction analyses of these proteins. Bacterially expressed, purified recombinant lamin A and LAP2α in urea-containing buffer, were dialyzed alone or together into lamin assembly buffer containing 300 mM NaCl, a salt concentration where lamin A does not form higher assembly structures (Foeger et al., 2006), and analyzed the protein samples by sucrose gradient centrifugation. (Fig. 5A). While both proteins alone sedimented mostly to fractions 2 and 3 of the sucrose gradient, their sedimentation pattern shifted towards lower fractions in the protein mix, consistent with the formation of larger hetero-complexes (Fig. 5A). As a control, a LAP2α mutant lacking the C-terminus that mediates interaction with A-type lamins (Dechat et al., 2000) did not alter the sedimentation pattern of lamin A (Fig. 5B).

**Figure 5.**
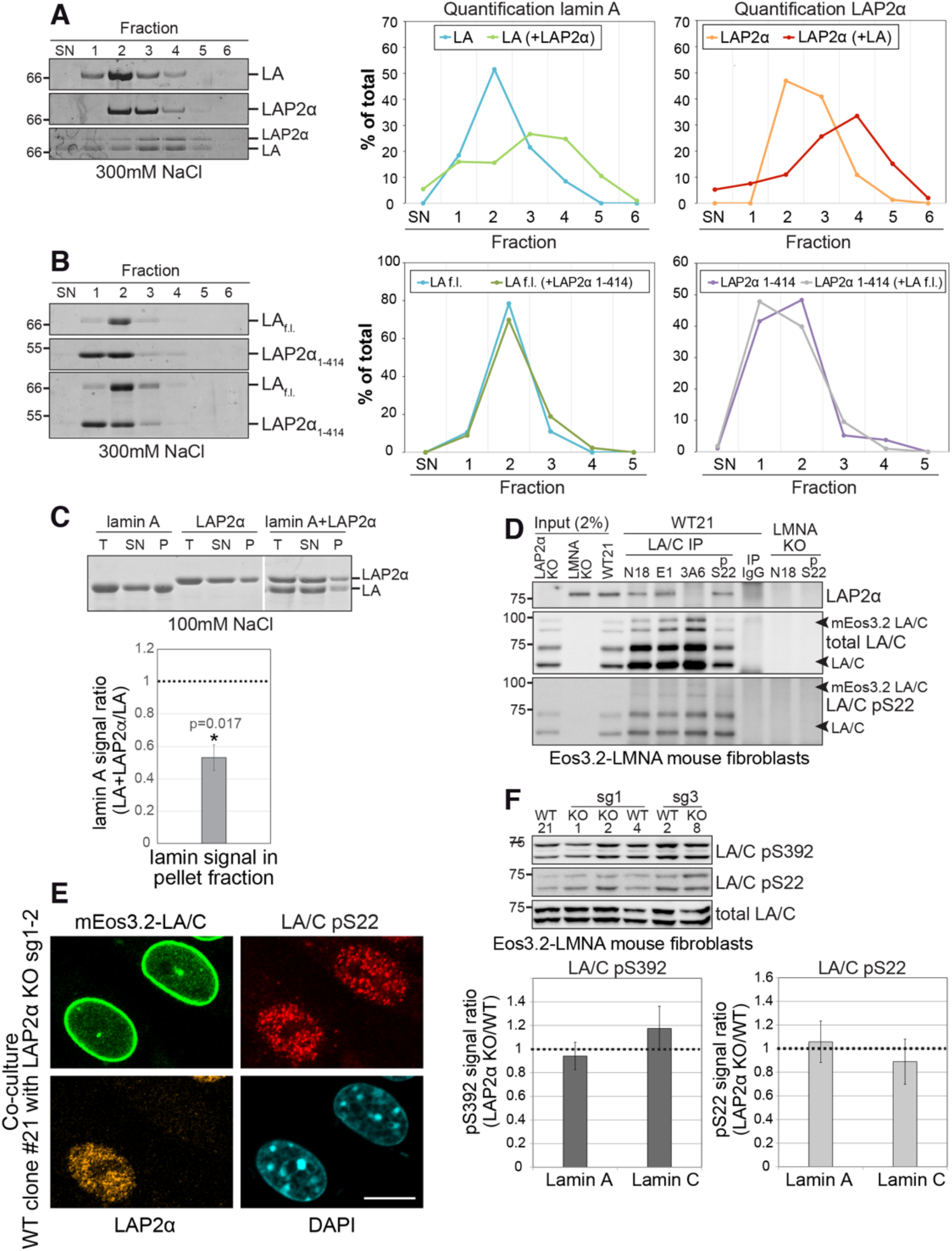
LAP2α binds to lamin A/C and inhibits their assembly without altering lamin phosphorylation. **(A, B)** Purified recombinant lamin A and LAP2α full length (A) or lamin A binding mutant LAP2α_1-414_ (B) were dialyzed either alone or together into assembly buffer with 300 mM NaCl. Samples were separated on a 10% to 30% sucrose gradient, followed by collection of fractions and quantification of protein bands. Exemplary Coomassie stained gels of fractions „supernatant” (SN) to 6 are shown on the left. The calculated protein amount per fraction (% of total protein) was plotted as curve chart on the right. **(C)** Lamin A and LAP2α were dialyzed either alone or together into assembly buffer as in (D), but with 100 mM NaCl, enabling formation of higher assembly lamin structures. After centrifugation, total (T), supernatant (SN) and pellet (P) fractions were analyzed on a Coomassie gel, protein bands were quantified and normalized to total protein levels. Graph displays lamin A levels in the pellet fraction of mixed samples (lamin A + LAP2α) as average fold difference ± S.E.M over samples with lamin A alone, n=5. **(D)** Lamin A/C was immunoprecipitated from WT clone #21 cells and LAP2α and lamin A/C knockout controls using the indicated antibodies recognizing different fractions of lamin A/C. Immunoprecipitates were analyzed by Western blotting using the indicated antibodies (anti lamin A/C 3A6, anti pS22 lamin A/C, anti LAP2α 1H11). **(E)** mEos3.2-*Lmna* WT clone #21 and LAP2α KO sg1-2 cells were co-cultured and processed for immunofluorescence microscopy using antibodies specific to lamins A/C phosphorylated at serine 22 (LA/C pS22), antibody 1H11 against LAP2α, and DAPI to visualize DNA. Bar: 10 μm. **(F)** WT clone #21 and isogenic LAP2α KO or WT clones were processed for Western blotting using antibodies against lamin A/C phosphorylated at specific residues as indicated or a pan-lamin A/C antibody (E1). Western blot signals for phosphorylated lamins were quantified, normalized to total lamins A and C and expressed as fold difference to the WT samples (graphs on the right). Graphs display average fold difference ± S.E.M. n_WT_=3, n_KO_=3.

Next, the proteins were dialyzed into buffer with a lower salt concentration (100 mM NaCl) allowing formation of higher assembly lamin structures (Foeger et al., 2006). High molecular mass lamin A assemblies were separated from those of lower oligomeric states by centrifugation and analyzed by gel electrophoresis (Fig. 5C). Strikingly, the presence of LAP2α reduced the amount of pelleted high molecular mass lamin A assemblies by half, indicating that LAP2α impairs lamin A assembly into larger structures (Fig. 5C). Thus, binding of LAP2α to lamin A may directly impair the formation of higher order lamin assemblies.

To confirm that LAP2α and lamin A/C interact *in vivo* in mEos3.2-lamin A/C cell lines, we performed co-immunoprecipitation using different lamin A/C antibodies. In accordance with previous data (Dechat et al., 2000; Gesson et al., 2016), antibodies to the lamin A/C N-terminus co-precipitated LAP2α from cell extracts (Fig. 5D, antibodies N18 and E1), whereas an antibody to the lamin A/C C-terminus failed to co-precipitate LAP2α (Fig. 5D, antibody 3A6), likely because binding of LAP2α to the lamin A/C C-terminus masks the epitope recognized by this antibody (Gesson et a., 2016). Thus, LAP2α interacts with lamin A/C *in vivo* and may thereby reduce the formation of higher order lamin assemblies in the nuclear interior.

Phosphorylation is another process potentially regulating lamin assembly and mobility, as particularly lamin phosphorylation at serines at position 22 and 392 was described to regulate lamina disassembly during mitosis, as well as lamin A/C localization, mobility and solubility in interphase cells (Heald & McKeon, 1990; Kochin et al., 2014). Thus, we wanted to test whether LAP2α may affect lamin A/C assembly indirectly, by regulating its phosphorylation status. Antibodies specifically recognizing lamin A/C phosphorylated at S22 revealed a similar punctate nucleoplasmic staining with complete absence of peripheral lamina staining in both LAP2α wildtype and knockout cells (Fig. 5E and Fig. S6C) (Kochin et al., 2014) and total levels of pS22 lamin A/C and pS392 were unchanged in knockout versus wildtype cells (Fig. 5E and F, Fig. S6B and C). In addition, the relative level of detergent/salt-extractable pS22 lamins A and C were unchanged in LAP2α knockout versus wildtype cells, in stark contrast to the significantly reduced levels of extractable total lamin A/C (Fig. 4C). Thus, LAP2α does neither effect the levels nor properties of phosphorylated lamins A and C. However, LAP2α can bind to pS22 lamins A/C as shown by coprecipitation from cell lysates (Fig. 5D, antibody pS22). We concluded that LAP2α binds both un(der)phosphorylated and pS22 lamin A, but affects preferentially the assembly state of un(der)phosphorylated lamin A/C, keeping it in a mobile, lower assembly state in a phosphorylation-independent manner.

### Reduced lamin A/C mobility in the absence of LAP2α decreases chromatin diffusion

To address the relevance of mobile intranuclear lamin A structures for nuclear functions and the consequences upon its loss in LAP2α-deficient cells, we tested chromatin mobility, since intranuclear lamins and LAP2α were found to bind to chromatin in the nuclear interior (Bronshtein et al., 2015; Gesson et al., 2016). Chromatin shows anomalous sub-diffusion, which – unlike free (normal) diffusion – is a motion that is slowed by constraints, such as temporal binding to nucleoplasmic lamins (Bronshtein et al., 2015). Depletion of lamins A/C was shown to increase chromatin motion and to change the type of diffusion towards normal unrestricted diffusion (Bronshtein et al., 2015). To test how depletion of LAP2α affects chromatin motion, we expressed fluorescently labeled TRF1 (to detect telomeres) in mEos3.2-*Lmna* WT#21 and LAP2α knockout cells and analyzed telomere trajectories to determine the diffusion volume of telomeres (Bronshtein et al., 2015; Vivante, Brozgol, Bronshtein, Levi, & Garini, 2019). Strikingly, telomere motion was significantly reduced in the absence of LAP2α when compared to wildtype cells (Fig. 6, left panel), and this effect was completely dependent on the presence of lamin A/C, since additional depletion of lamin A/C in LAP2α knockout cells (*Lmna*/*Lap2*α double knockout, see Fig. S7) increased the telomere motion volume to levels observed in *Lmna* single knockout fibroblasts (Fig. 6, right panel). Thus, the absence of LAP2α and the resulting changes in nucleoplasmic lamin mobility towards a more stable form have direct functional consequences on chromatin diffusion in the nuclear interior, significantly reducing telomere motion.

**Figure 6.**
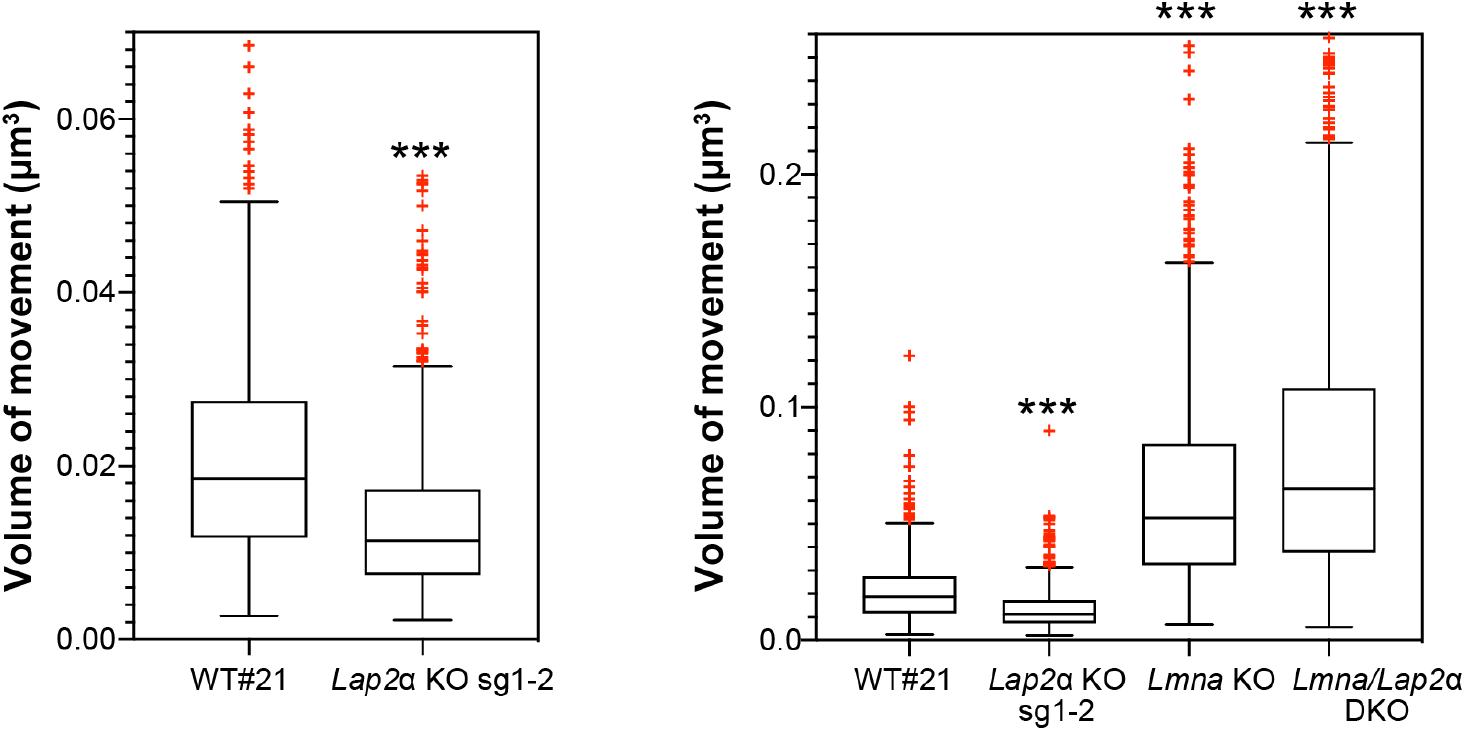
Lower lamin A/C mobility in the absence of LAP2α leads to a reduction in telomere movement. mEos3.2-*Lmna* WT clone #21 or mEos3.2-*Lmna* fibroblasts lacking LAP2α (sg1-2 *Lap2*α KO), lamin A/C (*Lmna* KO, see Fig. S7), or both (*Lmna*/*Lap2*α DKO) were transiently transfected with a plasmid expressing DsRed-TRF1 to fluorescently label telomeres. The volume of telomere motion was then calculated based on its trajectory. Box plots compare telomere motion between WT#21 and sg1-2 *Lap2*α KO (left; ***p < 0.0001 using Mann-Whitney U test) or between all 4 genotypes (right; Kruskal-Wallis test p < 0.0001; post tests for WT#21 ctrl versus each genotype are all highly significant with ***p < 0.0001). The median is depicted within the first and third quartiles; whiskers: minimal and maximal datapoint excluding outliers. n_WT#21_=471, n_*Lap2*αKO_=562, n_*Lmna*KO_=925, n_DKO_=1298.

In summary, our data provide novel insight into the formation and regulation of nucleoplasmic lamin A/C by LAP2α (Fig. 7). While the nucleoplasmic pool of lamin A/C can be formed independently of LAP2α, the properties of lamin A/C in the nuclear interior are significantly affected by LAP2α in a lamin serine 22 phosphorylation-independent manner. Binding of LAP2α impairs formation of higher order structures of lamin A/C and keeps them in a mobile and low assembly state, which in turn affects chromatin mobility.

**Figure 7.**
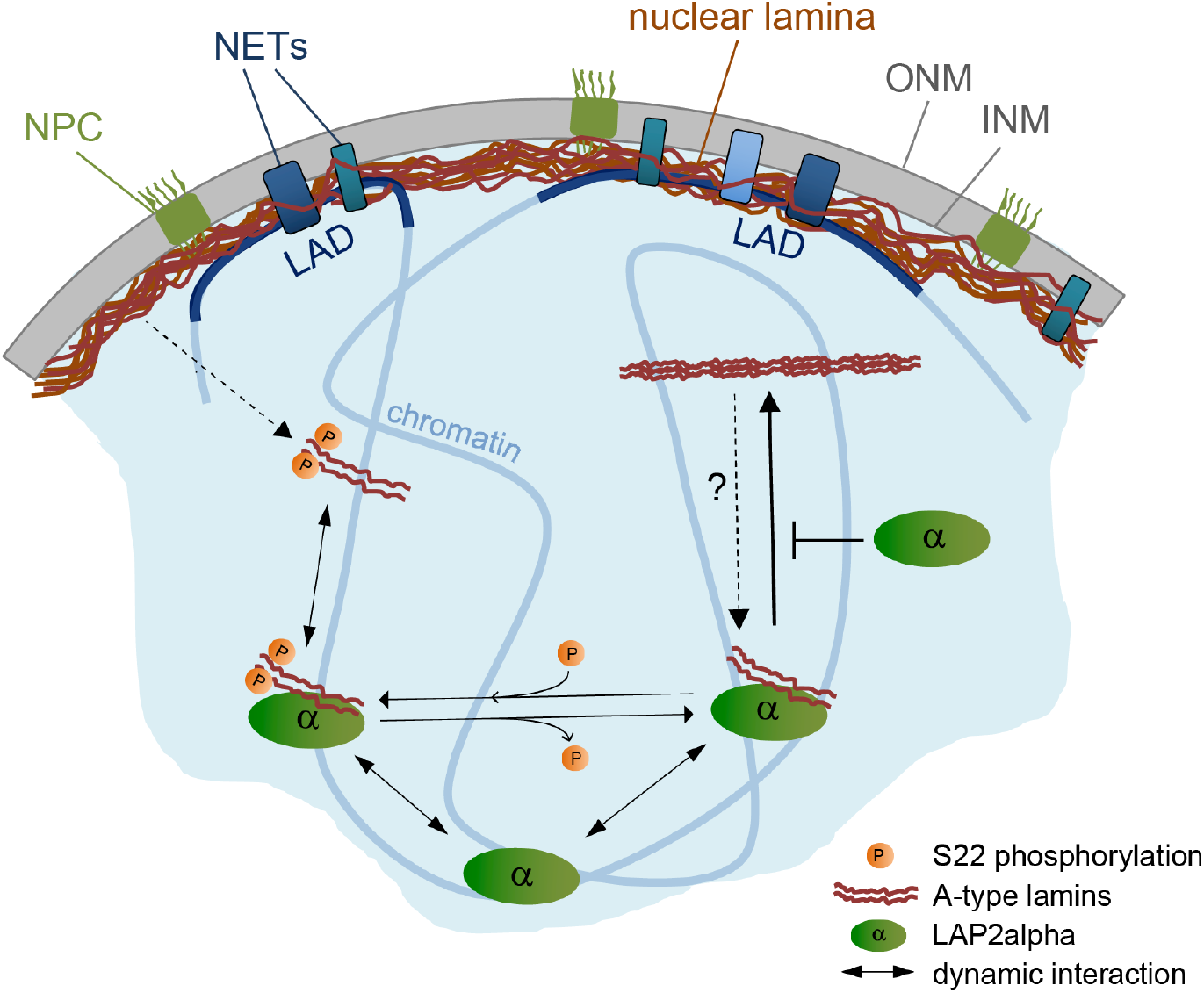
Model of LAP2α-dependent regulation of A-type lamins in the nucleoplasm. LAP2α dynamically interacts with lamins A and C in the nucleoplasm, independent of their phosphorylation status. While lamins at the periphery interact mainly with heterochromatic lamina-associated domains (LADs), nucleoplasmic A-type lamins and LAP2α bind to euchromatic genome regions regulating chromatin mobility. When hypophosphorylated lamins are not bound to LAP2α, they form larger, stable assemblies in the nucleoplasm. In the absence of LAP2α, these lamin A/C structures become more frequent, leading to altered, putatively less dynamic lamin A/C chromatin interaction and slower chromatin movement.

## Discussion

Here we show that LAP2α, a known binding partner of nucleoplasmic lamin A/C (Dechat et al., 2000; Naetar & Foisner, 2009) is essential to maintain lamins A and C in the nuclear interior in a mobile and low assembly state. This occurs in a lamin A/C S22 phosphorylation-independent manner, likely by inhibiting lamin assembly through direct interaction of LAP2α with lamin A/C.

Our study confirms and extends the finding that lamins A and C in the nuclear interior have fundamentally different properties compared to their counterparts at the nuclear periphery. While lamins at the nuclear periphery form stable 3.5 nm-wide filaments (de Leeuw et al., 2018; Moir et al., 2000; Shimi et al., 2008; Turgay et al., 2017), lamins in the nuclear interior are highly mobile (Broers et al., 1999; Bronshtein et al., 2015; Shimi et al., 2008) and easily extractable in salt and detergent-containing buffers (Kolb et al., 2011; Naetar et al., 2008), suggesting that they comprise mostly dimers and short polymers. However, very little is known about the mechanisms securing these unique properties of lamins in the nuclear interior, given that in solution they have a strong drive to assemble and tend to form higher-order structures (Aebi, Cohn, Buhle, & Gerace, 1986; Moir, Donaldson, & Stewart, 1991). Immunofluorescence microscopy has previously shown that intranuclear lamin A/C staining is significantly reduced in cells and tissues from LAP2α knockout mice (Naetar et al., 2008), leading to a model, where LAP2α is required for the formation and/or maintenance of the nucleoplasmic lamin pool (Naetar & Foisner, 2009). Based on data shown in this study, we propose a modified and updated version of the model suggesting that LAP2α is essential to maintain the highly mobile and soluble state of lamins in the nuclear interior. In the absence of LAP2α, lamins in the nucleoplasm are not lost but they are transformed into more stable, immobile structures (Fig. 7) that can no longer be recognized by lamin A/C antibodies directed to an N-terminal epitope, which – in wildtype cells – favors nucleoplasmic over peripheral lamin staining (Gesson et al., 2016). These findings can be explained by the formation of higher-order lamin structures and/or lamin complexes in the nuclear interior, whose properties may resemble more those of lamins at the nuclear periphery. This is supported by the increased resistance of nucleoplasmic lamins to extraction with detergent and salt-containing buffers and their decreased mobility in the absence of LAP2α. Moreover, high resolution microscopy revealed significantly more and, importantly, larger extraction-resistant structures in LAP2α knockout versus wildtype cells. As for potential mechanisms describing how LAP2α can maintain lamins in a soluble and dynamic state, our *in vitro* data using purified recombinant LAP2α and lamin A protein suggest that direct binding of LAP2α to lamins may prevent them from assembling into higher-order structures. Alternatively, besides affecting lamin assembly LAP2α may regulate the interaction of A-type lamins with chromatin. In this model, the absence of LAP2α would lead to a tighter and more stable association of lamins with chromatin, which would in turn also lead to an increased extraction resistance and possibly decreased mobility. In support of this model, the post-fixation digestion of formaldehyde-fixed cells with DNase/RNase fully recovered the intranuclear lamin A/C staining using antibodies to the lamin A/C N-terminus, and chromatin motion was slowed down upon LAP2α knockout in a lamin A/C dependent manner. However, the finding that the intranuclear lamin structures are resistant to 500 mM high salt, which is expected to extract chromatin-associated proteins (Herrmann, Avgousti, & Weitzman, 2017), supports the notion that larger lamin A structures can be formed independently of chromatin. As these different scenarios are not mutually exclusive, we favor a model in which the absence of LAP2α induces the formation of higher-order lamin A assemblies, which in turn may bind more tightly and stably to chromatin (Fig. 7).

As phosphorylation of lamins provides another means to regulate lamin filament (dis)assembly in mitosis (Heald & McKeon, 1990; Ward & Kirschner, 1990) and to influence their assembly state, solubility and mobility in interphase (Buxboim et al., 2014; Kochin et al., 2014; Moiseeva, Lopes-Paciencia, Huot, Lessard, & Ferbeyre, 2016), we tested the relationship between lamin phosphorylation and interaction with LAP2α. We found that LAP2α loss does neither affect total levels of lamins phosphorylated at serines 22 and 392 nor does it change the localization and solubility of phosphorylated lamins. Vice versa, LAP2α can interact with both phosphorylated and un(der)phosphorylated lamins, but it affects only the solubility and assembly state of un(der)phosphorylated lamins. Thus, interaction with LAP2α and lamin phosphorylation seem to be two independent, non mutually exclusive mechanisms for keeping lamins in a low assembly state.

Surprisingly, the initial formation of the nucleoplasmic lamin A/C pool does not require LAP2α, as it can form normally from newly synthesized pre-lamin A and from soluble mitotic lamins in LAP2α knockout cells. Thus, we propose that these processes may primarily depend on other mechanisms, such as lamin A/C phosphorylation. At the onset of mitosis, lamins are targeted by cyclin-dependent kinase 1 (Cdk1) particularly at serines 22 and 392 in the N-terminal head and proximal to the C-terminal end of the rod, respectively (Ward & Kirschner, 1990), leading to a complete disassembly of the lamina (Dechat et al., 2004; Heald & McKeon, 1990; Moir et al., 2000). During post-mitotic nuclear envelope reassembly, lamins A and C are imported into the nucleoplasm of newly formed nuclei, followed by gradual dephosphorylation and re-assembly at the nuclear periphery (Moir et al., 2000; Steen, Beullens, Landsverk, Bollen, & Collas, 2003; Steen & Collas, 2001). It is tempting to speculate that the majority of lamins A and C is still phosphorylated at S22 and possibly other residues when they are re-imported into the nucleoplasm after post-mitotic nuclear membrane reassembly, preventing their incorporation into the peripheral lamina. In the nucleoplasm, a fraction of the phosphorylated lamins may then bind to LAP2α. When lamins are gradually dephosphorylated during G1 (Steen et al., 2003; Steen & Collas, 2001), LAP2α-bound lamin A/C may be maintained in a low assembly state, while free lamin A/C can assemble into the lamina at the periphery. Thus, in the absence of LAP2α, the nucleoplasmic lamin A/C pool may still form, but, whereas phosphorylated lamins stay soluble independently of LAP2α, hypophosphorylated lamins tend to form higher-order structures at the “wrong” position, i.e. within the nucleoplasm as opposed to the nuclear periphery in LAP2α knockout cells. This model suggests that in wildtype cells, there are 3 pools of intranuclear A-type lamins (Fig. 7): 1) lamins phosphorylated at S22 (and possibly other residues) that are soluble independent of LAP2α; 2) hypophosphorylated lamins bound to LAP2α that are soluble independent of their phosphorylation status; and 3) hypophosphorylated lamins not bound to LAP2α that tend to form immobile and extraction-resistant structures in the nuclear interior. Previous reports and our own data showing that a small fraction of nucleoplasmic A-type lamins displayed low mobility (Bronshtein et al., 2015; Shimi et al., 2008) in wildtype cells supports this model. In the absence of LAP2α, the balance between these different intranuclear lamin structures is disturbed towards increased levels of immobile, higher order lamin A/C structures.

What is the physiological relevance of a dynamic lamin A/C pool in the nucleoplasm? We propose that these unique dynamic properties allow intranuclear A-type lamins to fulfill a set of functions that is different from those covered by the peripheral lamina. One of the most prominent examples for different functions of the peripheral versus intranuclear lamins is their different role in chromatin organization. While the peripheral lamina is well known to mediate stable interaction with heterochromatic genomic regions, termed lamina associated domains (LADs) (Kind et al., 2013; van Steensel & Belmont, 2017), thereby contributing to gene silencing in general (Leemans et al., 2019) or during differentiation (Robson et al., 2016), lamins in the nuclear interior have a much more complex role in regulating open chromatin and gene expression. Lamin A/C in the nuclear interior interact primarily with open euchromatic regions of the genome (Gesson et al., 2016; E. G. Lund, Duband-Goulet, Oldenburg, Buendia, & Collas, 2015) and affect their epigenetic landscape (Bianchi et al., 2020; Briand et al., 2018; Cesarini et al., 2015; Oldenburg et al., 2017). Furthermore, lamin A/C has also been shown to directly interact with promoters and enhancers and seems to be involved in both positive and negative gene regulation (Ikegami et al., 2020; E. Lund & Collas, 2013; E. Lund et al., 2013). It is conceivable that gene regulatory pathways have to be highly flexible and dynamic to respond efficiently to internal and external cues and thus require a dynamic lamin complex that can dynamically associate with and regulate chromatin. Indeed, loss of LAP2α (Gesson et al., 2016) or disease-linked mutations in lamins (Bianchi et al., 2020; Briand et al., 2018; Ikegami et al., 2020; Oldenburg et al., 2017) affect epigenetic regulation and/or gene expression.

Another function of nucleoplasmic lamins is to regulate the mobility of chromatin in the 3-dimensional nuclear space. Dynamic interactions of lamin complexes with telomeres and other genomic loci has been shown to limit free diffusion of chromatin inside the nucleus (Bronshtein et al., 2015). We demonstrate here that knockout of LAP2α reduces telomere motion in a strictly lamin A/C-dependent manner, most likely through the formation of more stable lamin A/C structures that bind chromatin more tightly and stably. It is reasonable to assume that a tightly regulated movement of genes inside the nucleus is important for controlling coordinated regulation of gene expression during various cellular processes.

Overall, we propose that lamin A/ C complexes are in a dynamic state, established by LAP2α, which is a prerequisite for their multiple roles in chromatin regulation both on a genome wide scale and in a gene specific manner.

## Materials and methods

### Generation and cultivation of cell lines

To generate mEos3.2-*Lmna* mouse dermal fibroblast cell lines, immortalized fibroblasts derived from the back skin of wildtype and LAP2α knockout littermates (Naetar et al., 2008) were transfected with the vector pSpCas9(BB)-2A-GFP (pX458, plasmid #48138 from Addgene, Watertown, MA) (Ran et al., 2013) carrying m*Lmna*-specific sgRNA1 (see ‘Primer and sgRNA sequences’) and Cas9 from *S. pyogenes* with 2A-EGFP, and a *Lmna*-specific targeting construct, where the fluorophore mEos3.2 (Zhang et al., 2012) was inserted in frame into exon 1 of the *Lmna* gene in front of the first codon (Fig. 3A, see also ‘Generation of *Lmna*-specific targeting construct’). Cells were sorted for EGFP expression using a FACS Aria Illu (Becton Dickinson, Franklin Lakes, NJ), cultivated for 14 days, followed by a single cell sort for mEos3.2-expressing cells. Single cell clones were outgrown and further characterized by genotyping PCR, long-range PCR, sequencing of the targeted locus and Western blotting.

The wildtype mEos3.2-*Lmna* clone #21 was selected to create isogenic *Lap2*α knockout clones, as well as *Lmna* knockout and *Lmna*/*Lap2*α double knockout clones. Cells were transfected with the vector pSpCas9(BB)-2A-mCherry (modified pSpCas9(BB)-2A-GFP, see also ‘Vectors’) carrying m*Lap2*α-specific sgRNA1 or 3, or m*Lmna*-specific sgRNA2 (see ‘Primer and sgRNA sequences’) and Cas 9 from *S. pyogenes* with 2A-mCherry. mCherry-positive single cells were FACS-sorted and knockout clones were identified by Western blot. Knockout clones and wildtype control clones were further characterized by sequencing of a PCR product derived from isolated genomic DNA spanning the expected Cas9 cut site (primers m*Lap2*α-f and -r, or m*Lmna*-f and -r: see ‘Primer and sgRNA sequences’). Sequences were analyzed using the TIDE software available online (https://tide.nki.nl/) (Brinkman, Chen, Amendola, & van Steensel, 2014). Absence of continuous Cas9 expression was verified by mCherry FACS analysis using an LSR Fortessa (Becton Dickinson).

To generate LAP2α knockout HeLa cell clones, HeLa cells were transfected with pSpCas9(BB)-2A-GFP carrying h*LAP2*α-specific sgRNA1 or 2 (see ‘Primer and sgRNA sequences’). GFP-positive cells were sorted and plated for generation of single cell clones. LAP2α knockout clones were identified by Western blotting. Knockout clones and wildtype control clones were further characterized by sequencing of a PCR product spanning the expected Cas9 cut site (primers h*LAP2*α-f and -r: see ‘Primer and sgRNA sequences’). Sequences were analyzed using TIDE. Absence of continuous Cas9 expression was verified by GFP FACS analysis.

All cells were maintained at 37°C and 5% CO_2_ in Dulbecco’s modified Eagle’s medium (DMEM) supplemented with 10% fetal calf serum (FCS), 2 mM glutamine, 50 U/ml penicillin and 50 μg/ml streptomycin (P/S) (all from Sigma-Aldrich, St. Louis, MO). Non-essential amino acids (PAN-Biotech, Aidenbach, Germany) were routinely added for mouse dermal fibroblasts. After sorting, cells were kept in medium additionally containing Antimycotic-Antibiotic (Gibco/Thermo Fisher Scientific, Waltham, MA) and Plasmocin (25 μg/ml, Invivogen, San Diego, CA) to prevent sorter-induced contamination.

### Live cell imaging

Cells were plated on 35mm glass-bottom μ-dishes (Ibidi, Gräfelfing, Germany) in high-glucose phenol-red free DMEM (Gibco Fluorobrite) supplemented with 10% FCS, L-glutamine, and P/S. For mEos3.2 cells, dishes were pre-coated with Cell-tak (Corning, Corning, NY) according to manufacturer’s instructions. HeLa cells were transfected with plasmids expressing EGFP-tagged pre-lamin A, mature lamin A, pre-lamin A delK32 or mature lamin A delK32 (see’Vectors’) using nanofectin (PAA Laboratories, Toronto, Canada) according to manufacturer’s instructions. To avoid substantial overexpression, a maximum of 1μg plasmid DNA per 35mm dish was used. Cells were imaged 5 hours or 24 hours post transfection.

Imaging was performed using a Visitron Spinning disc microscope (Zeiss, Oberkochen, Germany) under controlled environmental conditions at 37°C and 5% CO_2_, using a Plan-Apochromat 63x/1.4 Oil DIC III objective and an EM-CCD camera. Images (Z stacks) were obtained by automated multipositioning (40 positions per 35mm dish) every 20 minutes with the same excitation strength and exposure time using Visiview 4.4 software (Visitron Systems, Puchheim, Germany). Images were processed in FIJI. For experiments with HeLa cells, all time-lapse images were smoothed by 1-pixel radius Gaussian Blur filter, the frame was cropped to fit the cells of interest and substacks were created from Z stacks to include the focal plane of appropriate time points starting with the beginning of mitosis until late G1. These substacks were then used to create short movies and the images presented in the results section. The data were quantified manually by drawing a 3 pixel wide bar across the nuclei of selected cells and creating plot profiles for each cell. The peripheral (P) lamin signal was calculated by averaging two peak measurements, one from each end of the drawn line, and the nucleoplasmic (N) intensity as the average of 30 measurements spanning the center of the drawn line, followed by calculation of the N/P ratio.

Automated quantification of the N/P ratio for mEos3.2-lamin A/C cells was done using custom-made FIJI plugins, where mitotic cells are manually extracted from the images, followed by automatic analysis of time series stacks and data visualization of nucleoplasmic to peripheral signal intensity ratio over time. Specifically, frames, slices and fields of view with relevant cells were manually selected and extracted for automated analysis. Nuclei were segmented by thresholding, marking regions of interest. These regions were further reduced and subtracted to define nucleoplasm and nuclear periphery. Average intensity values of mEos3.2 were extracted from these regions and the signal ratio (nucleoplasm/periphery) was calculated.

### Immunofluorescence and cell cycle staging using DAPI

For standard immunofluorescence (IF), cells were seeded on uncoated glass coverslips (1.5H, Marienfeld-Superior, Lauda-Königshofen, Germany) and fixed with 4% paraformaldehyde for 10 min at room temperature. To stop fixation and permeabilize the cells, coverslips were incubated in PBS with 0.1% Triton X-100 and 50 mM NH_4_Cl, followed by incubation in primary and secondary antibody (for a list of all antibodies see ‘Antibodies’). To reverse chromatin-dependent epitope masking of nucleoplasmic lamins, cells were treated with 100 μg/ml DNase I and 100 μg/ml RNase A for 30 minutes at room temperature. DNA was stained with DAPI (0.5 μg/ml) and cells were mounted using Vectashield (Vector Laboratories, Burlingame, CA). Immunofluorescence slides were routinely imaged using an LSM710 confocal microscope (Zeiss) equipped with a Plan-Apochromat 63x/1.4 Oil DIC M27 objective and standard photomultiplier tubes (PMTs) for sequential detection, as well as an Airyscan detector for high resolution imaging. Image acquisition was done using Zeiss ZEN 2.1 software, followed by image processing using FIJI, including adjustment of the digital offset for high resolution Airy scan images to avoid over- and undersaturation of the fluorescent signal. IF images with standard resolution were smoothened by 1-pixel radius Gaussian Blur filter. Extraction-resistant lamin A/C structures were quantified using a custom-made FIJI plugin, where cells were first defined using the FIJI built-in auto-threshold function on average Z projections of mEos3.2-lamin A/C, followed by identification and quantification (number and area) of intranuclear structures using the threshold function with the built-in “Huang” algorithm, followed by the particle analyzer function, including particles between 0.0001 and 5μm, but excluding a rim of 0.8 μm from the nuclear periphery to avoid counting of signals within the nuclear lamina.

For DAPI-based cell cycle staging and combined determination of nucleoplasmic to peripheral ratio of mEos3.2-tagged lamin A/C, cells were imaged using the Visitron Spinning disc microscope (Zeiss) and a custom-made slide scanner, allowing the automated acquisition of multiple image stacks (400 fields of view/sample). Images were analyzed using a custom-made FIJI plugin, where the cell cycle stage of each cell was determined based on the DAPI intensity and a cell cycle profile was created. Initial regions of interest were extracted by segmenting nuclei using thresholding and watersheding. DAPI intensity was measured by summing the integrated density of DAPI signals within regions of interest. Initial regions of interest were further reduced and subtracted to define nucleoplasm and nuclear periphery. Intensity of the nucleoplasm was measured by taking the median value of the minimum projection of the regions of interest. Intensity of the nuclear periphery was measured by taking the upper quartile of the maximum projection of the regions of interest. The nucleoplasmic/peripheral lamin A/C ratio of the cells was calculated and plotted against the cell cycle stage.

### Continuous photobleaching and fluorescence correlation spectroscopy

Continuous photobleaching was performed using a confocal microscope (Leica TCS SP8 SMD, Leica Microsystems, Wetzlar, Germany). For illumination, the PicoQuant Picosecond Pulsed Diode Laser Head of 40 MHz 470 nm was used (laser intensity was set to ~1 μW). The light is focused onto a small confocal volume through a 63x water immersion objective lens with NA=1.2 (Leica HC PL APO 63x/1.20 W CORR). The signal is detected by sensitive detectors (Leica HyD SMD) within the detection window (500-600 nm). A specific point was chosen in the nuclear interior and a “point measurement” was performed for measuring the intensity at a high frequency (~1 KHz) for approximately 60 seconds per cell. CP data analysis was performed with Matlab and a custom-made algorithm (Bronshtein et al., 2015).

Fluorescence correlation spectroscopy (FCS) curves were extracted from the region where the fluorescence intensity is constant (approximately 30 seconds after beginning of the measurement) in order to avoid inaccuracy due to bleaching. The FCS analysis was done by SymPho Time software (PicoQuant, Berlin, Germany). The best fit was achieved with the FCS Triplet 3D fitting model:

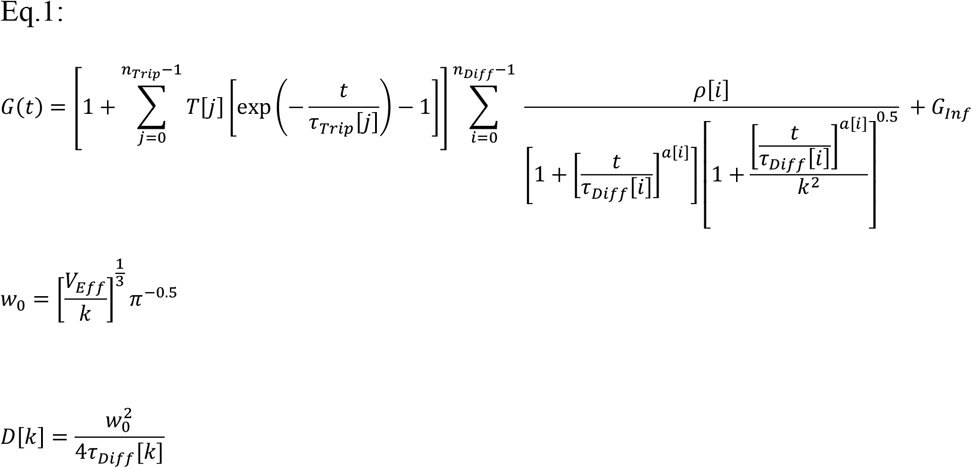

*Model parameters*:

Number of triplet (dark) states *n*_*Trip*_ = 1
Number of independently diffusing species *n*_*Diff*_ = 2
Effective excitation volume *V*_*Eff*_ = 0.267[*fl*]
Length to diameter ratio of the focal volume *k* = 7.92
Anomaly parameter of the i^th^ diffusing species *a*_1,2_ = 1
*Fitting parameters*:

Contribution of the i^th^ diffusing species - *ρ*[*i*]
Diffusion time of the i^th^ diffusing species - *τ*_*Diff*_[*i*]
Dark (triplet) fractions of molecules - *T*
Lifetime of the dark (triplet) states - *τ*_*Trip*_
Correlation offset - *G*_*Inf*_
Effective lateral focal radius at 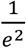 intensity - *w*_0_
Diffusion constant of the k^th^ diffusing species - *D*[*k*]

### Biochemical extraction experiments

Cells were washed twice with PBS (Sigma-Aldrich) and directly scraped off the plate in cold extraction buffer on ice (20 mM Tris-HCl pH7.5, 150 mM NaCl, 2 mM EGTA, 2 mM MgCl_2_, 0.5% NP-40, 1mM DTT, 1U/ml Benzonase from Novagen/EMD Millipore, Temecula, CA, 1x Complete Protease Inhibitor Cocktail from Sigma-Aldrich, 1x Phosphatase inhibitor cocktail 2 and 3 from Sigma-Aldrich). For mouse fibroblasts extra NaCl was added to a final concentration of 500 mM. Extracts were incubated for 10 min on ice and then centrifuged for 10 min at 4000 rpm in a Megafuge 1.0R (Haereus, Hanau, Germany). Pellets were resuspended in equal volumes of extraction buffer and sonicated with a Bandelin Sonopuls HD200 sonicator (Bandelin, Berlin, Germany; settings: MS73/D at 50% intensity) for 3 sec to solubilize insoluble material. Total, supernatant and pellet fractions were analyzed by immunoblotting.

For *in situ* extraction of mEos3.2-lamin A/C mouse fibroblasts, cells were grown on glass coverslips (1.5H, Marienfeld) that were pre-coated with Cell-tak (Corning) according to manufacturer’s instructions to avoid detachment of cells due to extraction. Cells were washed twice with PBS and incubated for 10 min in extraction buffer containing 500 mM NaCl (without benzonase), followed by fixation in 4% paraformaldehyde according to the routine IF protocol (see ‘Immunofluorescence and cell cycle staging using DAPI’). To amplify the mEos3.2 signal, cells were stained with an anti-mEos2 antibody (Badrilla, Leeds, UK) following the standard IF protocol.

### Recombinant protein expression and sedimentation assays

Recombinant full length human LAP2α and the truncation mutant LAP2α1-414 were expressed in E. coli BL21 (DE3) using the plasmid pET23a(+) (Vlcek, Just, Dechat, & Foisner, 1999), which adds a 6x histidine-tag to the C-terminus of the expressed proteins for further affinity purification. Protein expression was induced in bacterial cultures at an OD_600nm_ of 0.6 – 0.7 with 0.5 mM Isopropyl-β;-D-thiogalactopyranosid (IPTG) for 3 hours. Bacterial pellets were then resuspended in 10 mM Tris-HCl pH 8.0, 100 mM NaCl, and 1 mM DTT, followed by cell lysis in the presence of 1x protease inhibitor cocktail (Sigma Aldrich). 5 μg/ml DNase I and 10 μg/ml RNase A were added and inclusion bodies of the respective recombinant proteins were pelleted by centrifugation. Poly-histidine-tagged recombinant proteins were purified using Ni-NTA-Agarose (Qiagen, Hilden, Germany) beads according to manufacturer’s instructions and were analyzed by SDS-polyacrylamide gel electrophoresis (SDS-PAGE). Pure protein fractions were dialyzed twice against 8 M Urea, 10 mM Tris-HCl pH 8.0, 300 mM NaCl and 1 mM DTT using a cut-off of 12-14 kDa to remove Imidazole and ß-Mercaptoethanol. Purified recombinant human wild type lamin A was a kind gift of Prof. Harald Herrmann, DKFZ, Heidelberg, Germany.

For sucrose density gradient centrifugation, recombinant proteins were dialyzed against 10 mM Tris-HCl pH 7.5, 1 mM DTT, 300 mM NaCl and 10% sucrose and were layered on top of a sucrose gradient ranging from 10% to 30% atop a 70% sucrose cushion in a centrifuge tube. Samples were centrifuged for 20 h at 190,000 x g in a SW-40 Rotor (Beckman Optima L-70 or Beckman Optima L-80 XP, Beckman Coulter, Brea, CA) at 4 °C. Collection of the sedimented protein layers was done by pipetting off fractions from the top to the bottom of the gradient, starting with the applied dialyzed protein sample (supernatant). Aliquots of all gradient fractions were analyzed on an SDS-polyacrylamide gel stained with Gel CodeTM Blue Safe Protein Stain (Thermo Fisher Scientific) and density measurement of the protein bands was done with ImageJ software.

To induce lamin assembly, recombinant proteins were dialyzed stepwise against 10 mM Tris-HCl pH 7.5, 1 mM DTT containing decreasing salt concentrations, starting at 300 mM NaCl, followed by 200 mM NaCl, and ending at 100 mM NaCl. Each step was done at room temperature for 30 min. Samples were then analyzed by centrifugation at 13,000 rpm for 10 min (Eppendorf table top centrifuge, Eppendorf, Hamburg, Germany) at room temperature. Supernatant and pellet fractions were analyzed by SDS-PAGE as described above for sucrose gradient fractions.

### Immunoprecipitation and Immunoblotting

For co-immunoprecipitation (IP), cells were scraped off the plate in IP buffer containing 20 mM Tris-HCl pH7.5, 150 mM NaCl, 2 mM EGTA, 2 mM MgCl_2_, 0.5% NP-40, 1 mM DTT, 1 U/ml Benzonase (Novagen), 1x Complete Protease Inhibitor cocktail (Sigma-Aldrich) and Phosphatase inhibitor cocktail 2 and 3 (Sigma-Aldrich), and incubated 10 min on ice. The soluble fraction after centrifugation for 10 min at 4,000 rpm in a Megafuge 1.0R (Haereus) was used as input for the IP (1 mg/IP). Incubation with antibody (5 μg/IP) was done overnight at 4°C, followed by incubation with BSA-blocked proteinA/G dynabeads (Pierce/Thermo Fisher Scientific) for 4 hours at 4°C. Beads were washed 3 times in IP buffer without benzonase, followed by elution of complexes from the beads using SDS PAGE sample buffer. Samples were analyzed by SDS PAGE and immunoblotting.

Immunoblots were treated with primary antibodies overnight (see ‘Antibodies’) and with horse radish peroxidase-coupled secondary antibodies (Jackson ImmunoResearch, Westgrove, PA) for one hour at room temperature. Signal detection was done using SuperSignal West Pico plus chemiluminescent substrate (Pierce/Thermo Fisher Scientific) and the ChemiDoc Gel Imaging system (Bio-Rad, Hercules, CA). Image analysis and quantification was done using the Image Lab software (Bio-Rad).

### Telomere tracking

Cells were transfected with the DsRed-TRF1 plasmid (Bronshtein et al., 2015) one day before imaging. For imaging, cells were placed in an incubator (Tokai, Shizuoka-Ken, Japan) mounted on an inverted Olympus IX-81 fluorescence microscope coupled to a FV-1000 confocal set-up (Olympus, Tokyo, Japan) using a UPLSAPO 60X objective lens with a numerical aperture of 1.35. Each nucleus was measured 50 times in three dimensions (3D) for a total time of 17.5 min. Imaris image analysis software package (Bitplane, Zurich, Switzerland) was used for correcting the nucleus drift and rotation and for identifying the coordinates of labelled telomeres. Only telomeres that were tracked over all the 50 time points were considered for further data analysis. For calculating the volume covered by each telomere during its whole motion, we used the Convex hull algorithm using a custom-made Matlab code.

### Quantitative real time PCR for detecting untagged and mEos3.2-tagged lamin A/C mRNA levels

RNA was isolated from mEos3.2-lamin A/C WT#21 cells using the RNeasy mini plus kit (Qiagen) according to manufacturer’s instructions. cDNA was synthesized from 500 ng total RNA using the RevertAid First Strand cDNA synthesis kit (Thermo Fisher Scientific) and analyzed by qPCR using the KAPA SYBR Green 2x PCR master mix (Kapa Biosystems, Wilmington, MA) in an Eppendorf Realplex 2 Mastercycler according to manufacturer’s instructions. Primers specific for untagged and tagged lamins A/C (wt LAC-f and -r; mEos LA/C-f and -r – see ‘Primer and sgRNA sequences’) were used to generate PCR products that were gel-extracted and used as a template in real time PCR to generate a standard curve (DNA concentration versus threshold cycle). WT#21 cDNA was then analyzed by real time PCR using the same primers and DNA/RNA concentration of the specific template was calculated from the standard curve.

### Cloning of *Lmna*-specific targeting construct

The *Lmna*-specific targeting construct was assembled from 4 fragments that were amplified by PCR using the following primer pairs and templates (see also ‘Primer and sgRNA sequences’):

| **Fragment** | **f primer (5'- 3')** | **r primer (5'- 3')** | **Template** |
| --- | --- | --- | --- |
| 1 | m*Lmna*-Frag1-f | m*Lmna*-Frag1-r | mouse genomic DNA |
| 2 | mEos-Frag2-f | mEos-Frag2-r | mEos3.2-C1 (Addgene, plasmid #54550) |
| 3 | m*Lmna*-Frag3-f | m*Lmna*-Frag3-r | mouse genomic DNA |
| 4 | m*Lmna*-Frag4-f | m*Lmna*-Frag4-r | mouse genomic DNA |

Primers were designed to generate overlapping fragments that were assembled using the Gibson assembly master mix (New England Biolabs, Ipswich, MA) according to manufacturer’s instructions. The vector pUC18 (Norrander, Kempe, & Messing, 1983) was digested with SalI and EcoRI (New England Biolabs) creating vector ends overlapping with fragment 1 and 4, respectively, and added to the Gibson assembly reaction. The final *Lmna* targeting construct in pUC18 was sequence-verified and contained *Lmna* exon 1 and its flanking non-coding sequences, where mEos3.2 was inserted in frame into exon 1 before the first codon (removing the start codon). An additional EcoRI site was inserted directly after mEos3.2 and the recognition sequence of *Lmna*-specific sgRNA1 was altered within exon 1 (5’ CACTCGGATCACCCGcCTaC 3’, mutations are indicated in lowercase) to avoid recutting and potential creation of Indels in the modified allele.

### Genotyping PCR and long range (LR) PCR for modified mEos3.2-*Lmna* knock-in allele

To identify clones with a modified mEos3.2-*Lmna* knock-in allele, genomic DNA was isolated from single cell clones using the QuickExtract DNA extraction solution (Epicentre/Lucigen, Middleton, WI) according to manufacturer’s instructions and analyzed for the presence of the wildtype and knock-in *Lmna* allele by PCR using the GoTaq green master mix (Promega, Madison, WI) and the following primer pairs (see also ‘Primer and sgRNA sequences’):

| PCR | Forward primer (5'- 3') | Reverse primer (5'- 3') | Product |
| --- | --- | --- | --- |
| WT | m*Lmna*-WT-f | m*Lmna*-WT-r | 213 bp (KI: 894 bp) |
| Knock-in | m*Lmna*-KI-f | m*Lmna*-KI-r | 250 bp |

LR-PCR was performed in clones with at least one knock-in allele to verify correct integration of the construct at the 3-prime and 5-prime side with one primer of each pair outside the homology region of the *Lmna* targeting construct (see also Fig. S3). PCR reactions were set up using the Q5 DNA polymerase (New England Biolabs) according to manufacturer’s instructions and the following primer pairs:

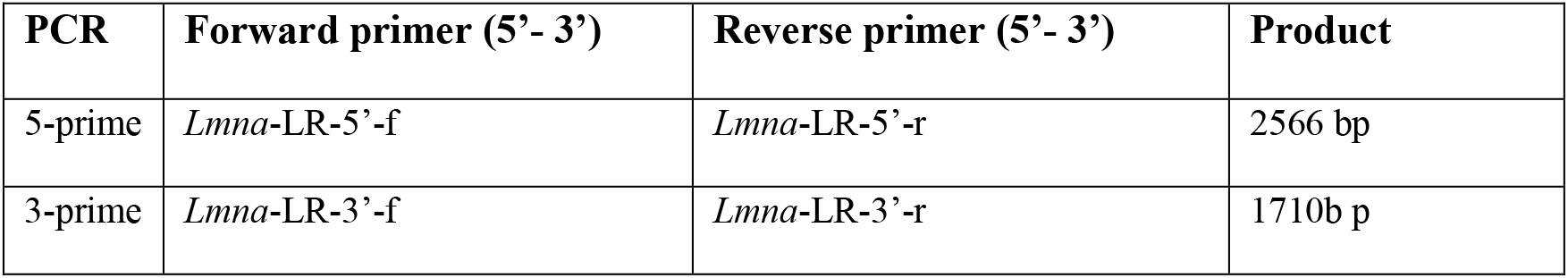

### Vectors

pSpCas9(BB)-2A-mCherry was generated from pSpCas9(BB)-2A-GFP (Addgene, plasmid #48138) by removing EGFP and the T2A sequence via EcoRI digestion. The final vector was then assembled using the Gibson assembly master mix (New England Biolabs) and 2 overlapping fragments containing the T2A sequence and mCherry generated by PCR using the following primers and templates (see ‘Primer and sgRNA sequences’):

| Fragment | Forward primer (5'- 3') | Reverse primer (5'- 3') | Template |
| --- | --- | --- | --- |
| T2A | T2A-f | T2A-r | pSpCas9(BB)-2A-GFP |
| mCherry | mCherry-f | mCherry-r | pLVX mCherry (Takara Bio) |

Vectors expressing different lamin A constructs were all derived from pEGFP-myc-LMNA (Moir et al., 2000). pEGFP-myc-Lamin A del K32, harboring a deletion of lysine at position 32 (Bertrand et al., 2012), was derived from pEGFP-myc-LMNA by site directed mutagenesis as previously described (Pilat et al., 2013). Wildtype and delK32 mature lamin A constructs were generated by PCR from pEGFP-myc-LMNA and pEGFP-myc-Lamin A delK32, respectively, using primers hLaminA-NheI-f and hLaminA-NheI-r.

### Antibodies

Antibodies used in this study were: rabbit antiserum to LAP2α (Vlcek, Korbei, & Foisner, 2002), monoclonal antibodies against LAP2α (1H11) or lamin A/C (3A6) (Max Perutz Labs monoclonal antibody facility, see also Gesson et al., 2016), hybridoma supernatant containing antibodies against the LAP2 common domain (Ab12) or the LAP2α-specific C-terminus (Ab 15-2) (Dechat et al., 1998), anti-lamin A/C N18 and E1 (both from Santa Cruz Biotechnology, Dallas, Texas), anti-lamin A/C phosphorylated at S22 (PA5-17113 from Invitrogen/Thermo Fisher Scientific or D2B2E from Cell Signaling, Danvers, MA) or S392 (ab58528, Abcam, Cambridge, UK), anti-lamin B1 (12987-1-AP, Proteintech, Rosemont, IL), anti-mEos2 (A010, Badrilla), anti-gamma-tubulin (T6557, Sigma-Aldrich), and anti-actin (I-19, Santa Cruz Biotechnology).

### Primer and sgRNA sequences

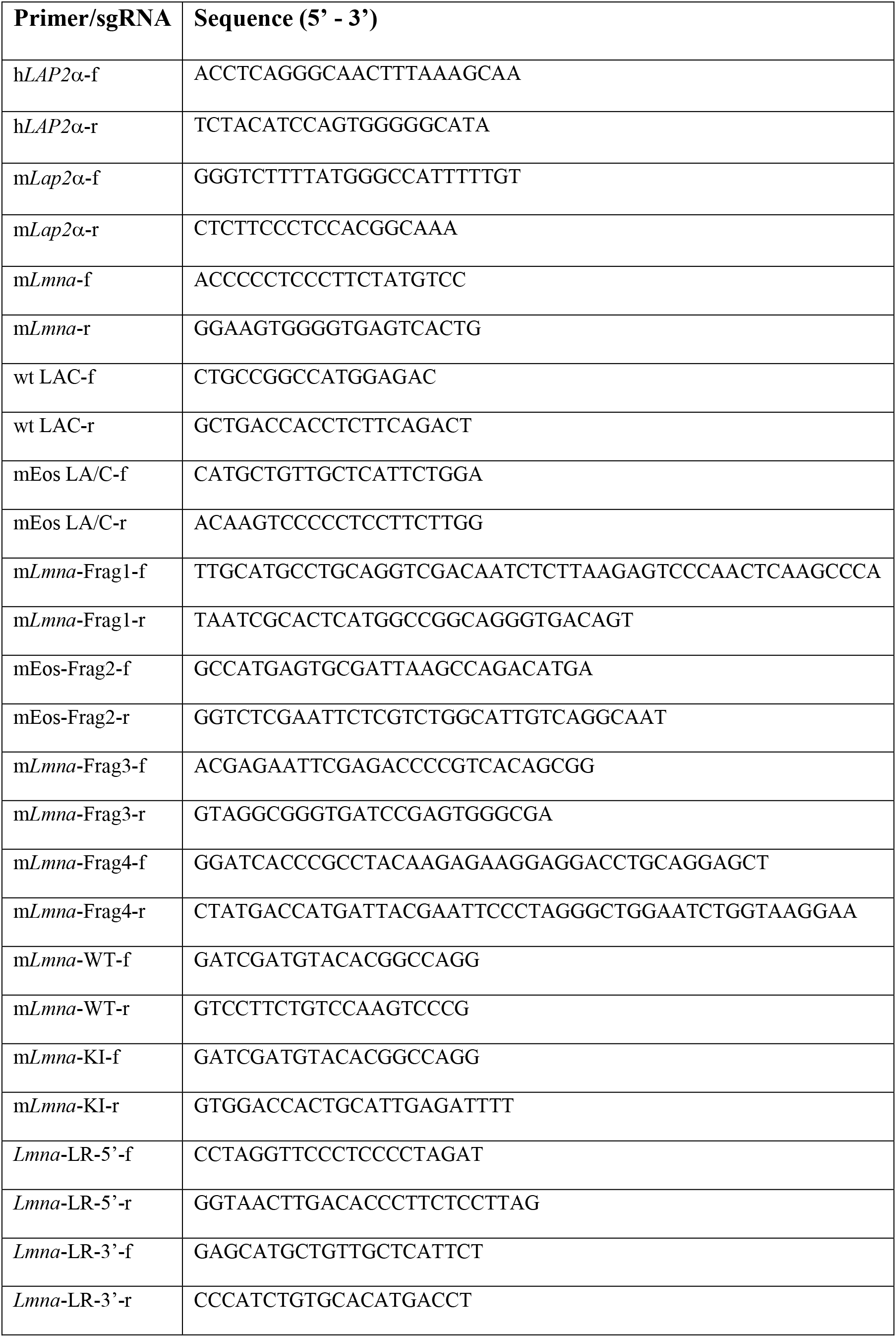
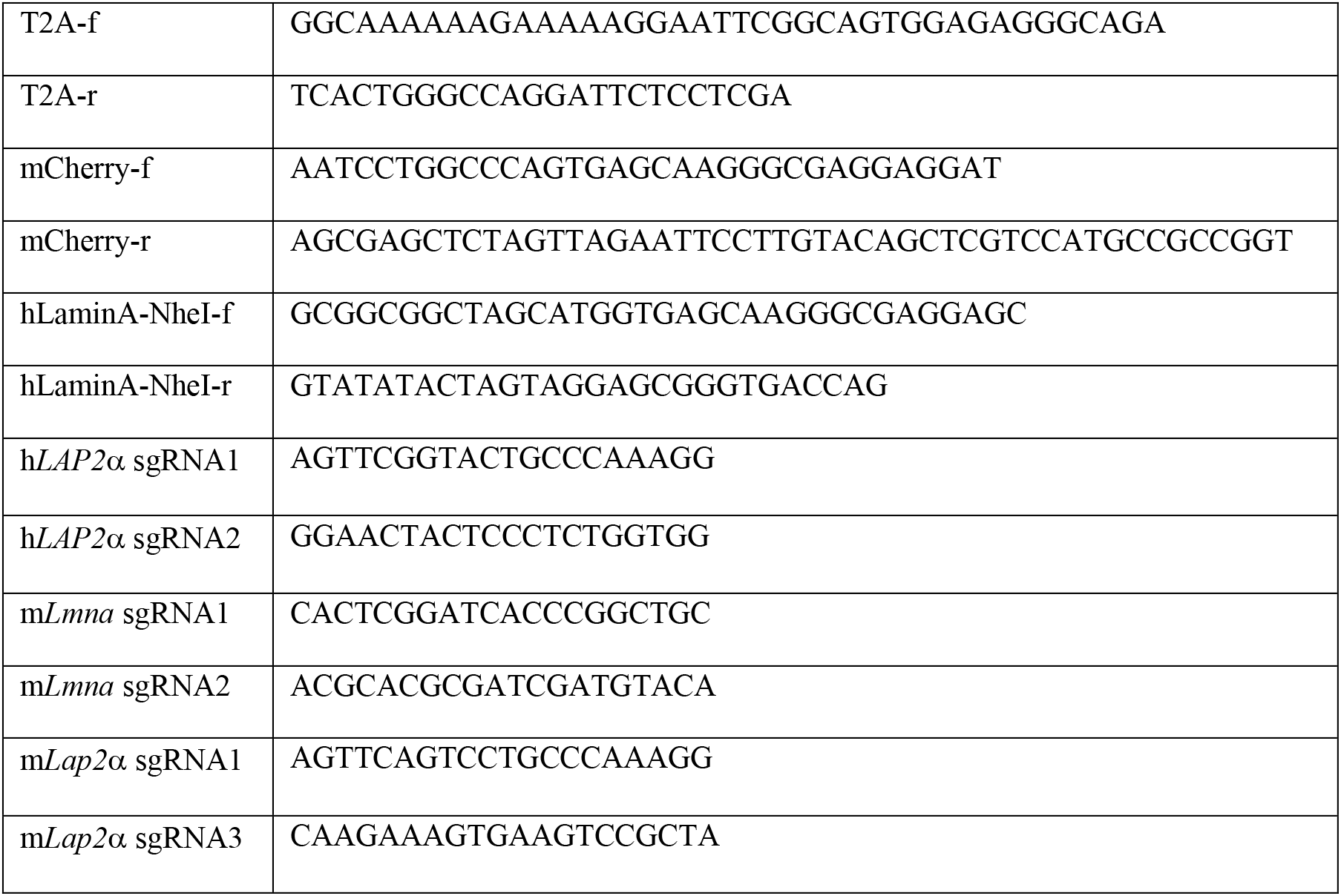

### Statistical analysis

To summarize experimental data sets, average, standard deviation and standard error of the mean were calculated and displayed in bar graphs. In specific cases, box plots were chosen to display data to better visualize data distributions. In box plots the median was depicted within the first and third quartiles with the whiskers representing minimal and maximal datapoints excluding statistical outliers (according to the method of Tukey using the inter-quartile range). For statistical analysis, normal distribution of data was tested using the D’Agostino-Pearson and Shapiro Wilk normality test. Additionally, quantile-quantile plots were created and visually inspected for normal distribution. For normally distributed data sets, the two-tailed student’s t test was used for statistical analysis (unpaired or paired, if data points were matched in pairs). The F test was used to determine whether variances of data sets are equal. Data expressed as proportions are not normally distributed and were transformed using the arcsin transformation (transformed 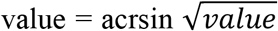). Ratios were transformed using the logarithmic transformation. If multiple comparisons were necessary (e.g. more than 2 data sets to compare with each other), ANOVA was used, including post-hoc tests for pairwise comparisons (Tukey). For non-normally distributed data, the Mann-Whitney U test was used, or, for multiple comparisons, the Kruskal-Wallis test.

## Acknowledgements

We are grateful to Harald Herrmann, DKFZ Heidelberg, Germany for the generous gift of recombinant lamin A and for his advice for lamin A *in vitro* assembly assays, and to Egon Ogris, Max Perutz Labs, Vienna, for the lamin A 3A6 antibody. We thank the Max Perutz Labs Biooptics facility for technical support with microscopic imaging and image analysis and technical support with FACS sorting.

## Additional Information

The authors declare that no competing interests exist.

## Funding

This study was funded by the Austrian Science Fund (FWF grant P26492-B20, P29713-B28 and P32512-B) to RF. KG is a recipient of a DOC Fellowship of the Austrian Academy of Sciences at the Max Perutz Labs, Medical University Vienna (ÖAW DOC 25725). NN was a recipient of an APART Fellowship of the Austrian Academy of Sciences at the Max Perutz Labs, Medical University Vienna (APART 11657). YG and IB would like to acknowledge financial support from the Israel Science Foundation (ISF) grant 1219/17 and from the S. Grosskopf grant for ‘Generalized dynamic measurements in live cells’. TD was a recipient of two COST Short Term Scientific Mission Fellowships (COST-STSM-BM1002-8698 and COST-STSM-BM1002-11436) and an EMBO short term fellowship (ASTF 316-2011).

## Author contributions

Conceptualization, N. Naetar, T. Dechat, Y. Garini and R. Foisner; Investigation, N. Naetar, K. Georgiou, C. Knapp, I. Brohnshtein, E. Zier, and P. Fichtinger; Formal analyses, N. Naetar, K. Georgiou, C. Knapp, I. Brohnshtein, and E. Zier; Methodology, C. Knapp and K. Georgiou; Resources, Y. Garini and R. Foisner; Project recruitment and administration, R. Foisner; Writing original draft, N. Naetar, K. Georgiou and I. Bronshtein; Writing, Review and editing, all; Supervision, N. Naetar, T. Dechat, Y. Garini and R. Foisner, Funding Acquisition, R. Foisner, N Naetar.

## Online supplemental material

Fig. S1 shows the characterization of HeLa LAP2α knockout clones created by CRISPR-Cas9 used in Fig. 1. Videos 1-4 show live-cell imaging of HeLa wildtype and LAP2α knockout cells expressing GFP-pre-lamin A, imaged either 5 hours (Videos 1 and 2, corresponding to Figure 1A) or 24 hours post transfection (Videos 3 and 4, corresponding to Figure 1B). Fig. S2 shows live cell imaging of HeLa cells expressing GFP-pre-lamin A, mature GFP-lamin A, GFP-ΔK32 pre-lamin A or mature GFP-ΔK32 lamin A (associated with Fig. 1). Videos 5-7 correspond to panels 2-4 in Fig. S2A and videos 8-10 correspond to panels 2-4 in Fig. S2B. Fig. S3 shows the characterization of mEos3.2-lamin A/C mouse dermal fibroblasts used in Figs 2–6. Fig. S4 shows the characterization of isogenic mEos3.2-lamin A/C LAP2α knockout cell lines used in Figs 2–6. Fig. S5 shows immunofluorescence images of various LAP2α knockout and wildtype mEos3.2 cells with different antibodies (associated with Fig. 3). Fig. S6 shows lamin A/C extraction and lamin phosphorylation in wildtype and LAP2α knockout HeLa cells (associated with Figs 4 and 5). Fig. S7 shows characterization of *Lmna* knockout and *Lmna/Lap2*α double knockout mouse dermal fibroblasts used in Fig. 6.

**Figure S1.**
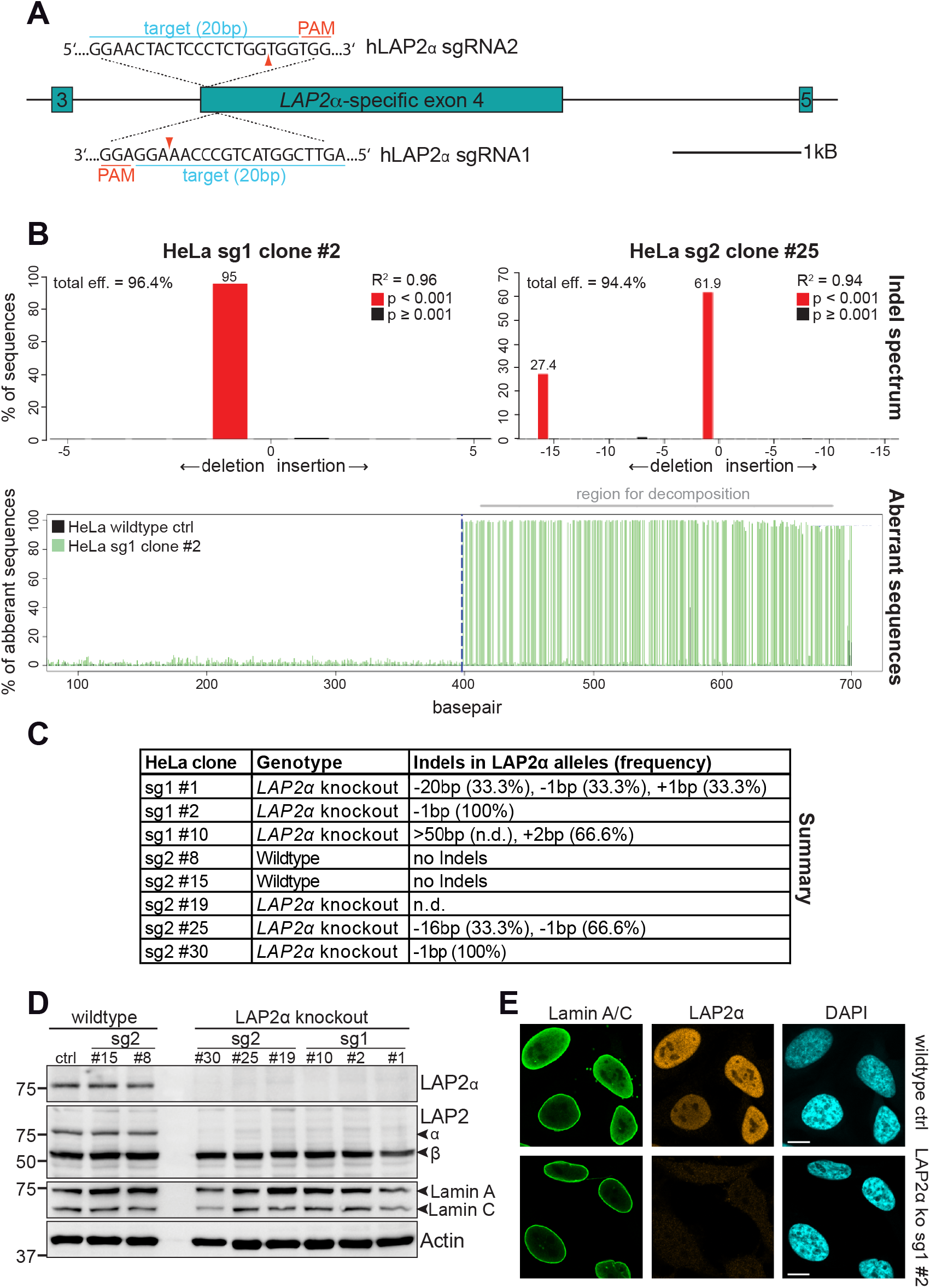
Generation of LAP2α knockout HeLa cell clones using CRISPR-Cas9. **(A)** Schematic view of exons 3-5 (bars) and adjacent introns (lines) of the human *LAP2* locus. Positions of the target sequences of LAP2α-specific sgRNAs 1 and 2 at the beginning of exon 4 are shown (blue). Protospacer-adjacent motif (PAM) sequences for each sgRNA are marked in red. Red arrowhead, expected Cas9 cut site. **(B)** A region spanning both cut sites was amplified from genomic DNA of the indicated clones and wildtype control cells by PCR. PCR products were sequenced and analyzed using the TIDE software (Brinkman et al., 2014). Upper graphs display detected Indels and their frequency in % (numbers atop red bars), lower graph shows an example alignment of wildtype and clone sg1 #2 sequences. Frequency of aberrant sequences, defined as sequences that differ from wildtype, are displayed on the Y axis and increase drastically after Indel-induced frame shifts at the expected cut site (dashed blue line). The region used for decomposition of individual sequences and Indel prediction is marked in grey. **(C)** Summary of generated HeLa clones and their detected Indels as determined by TIDE analysis. Allele frequencies were calculated based on TIDE results and clone ploidy. Alleles with deletions >50 bp cannot be analyzed by TIDE. **(D)** HeLa clones were processed for Western blot analysis using antibodies to the indicated antigens (anti lamin A/C 3A6, anti LAP2α Ab15-2). **(E)** HeLa control and LAP2α knockout sg1 #2 cells were processed for immunofluorescence microscopy using the indicated antibodies (anti lamin A/C 3A6, LAP2α rabbit antiserum) and analyzed using an LSM 710 confocal microscope. Bar: 10 μm.

**Figure S2.**
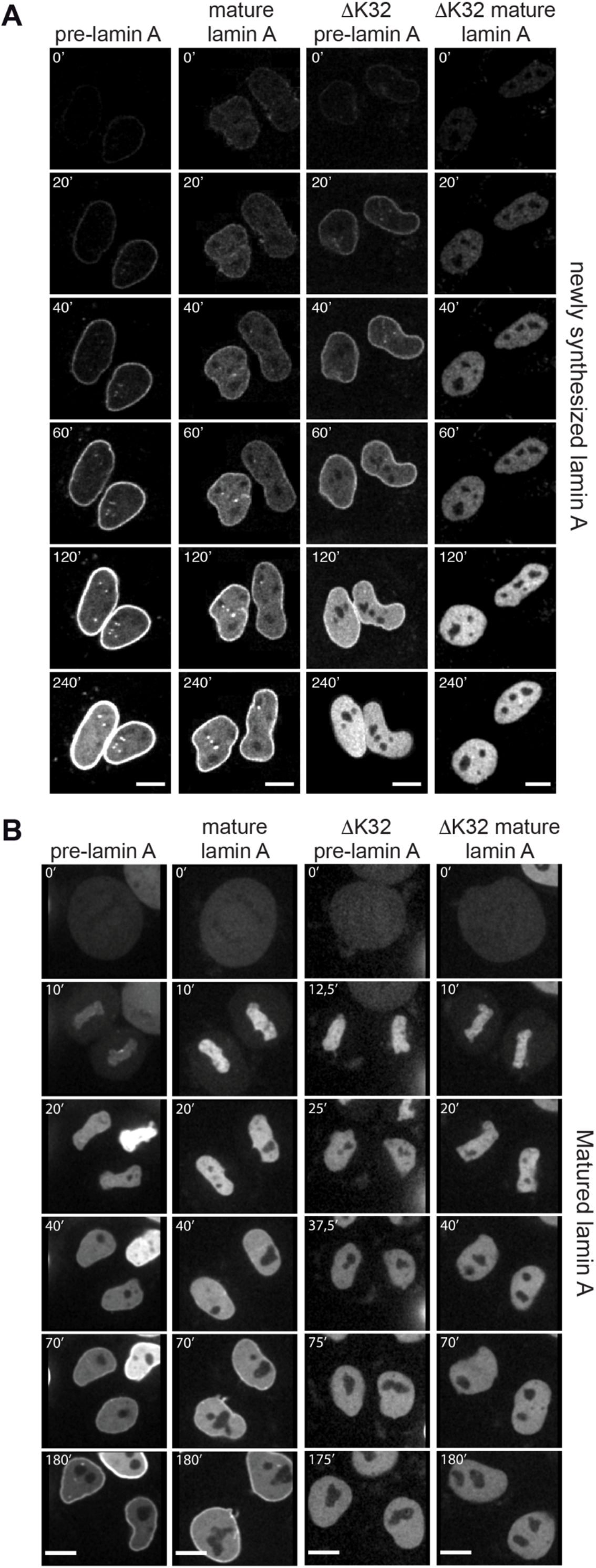
Newly expressed pre-lamin A and mitotically disassembled mature lamin A contribute to the nucleoplasmic lamin pool. HeLa cells were transiently transfected with either EGFP-pre-lamin A, EGFP-mature lamin A, EGFP-ΔK32 pre-lamin A or EGFP-ΔK32 mature lamin A as indicated, and analyzed by live-cell imaging 5 hours **(A)** or 24 hours **(B)** post-transfection. See also video files: Videos 5-7, corresponding to panels 2-4 in (A) and videos 7-10, corresponding to panels 2-4 in (B). For EGFP-pre-lamin A expressing HeLa cells imaged 5 and 24 hours post-transfection, see Video 1 and 3, respectively, associated with Figure 1. Numbers indicate minutes. Bar: 10 μm.

**Figure S3.**
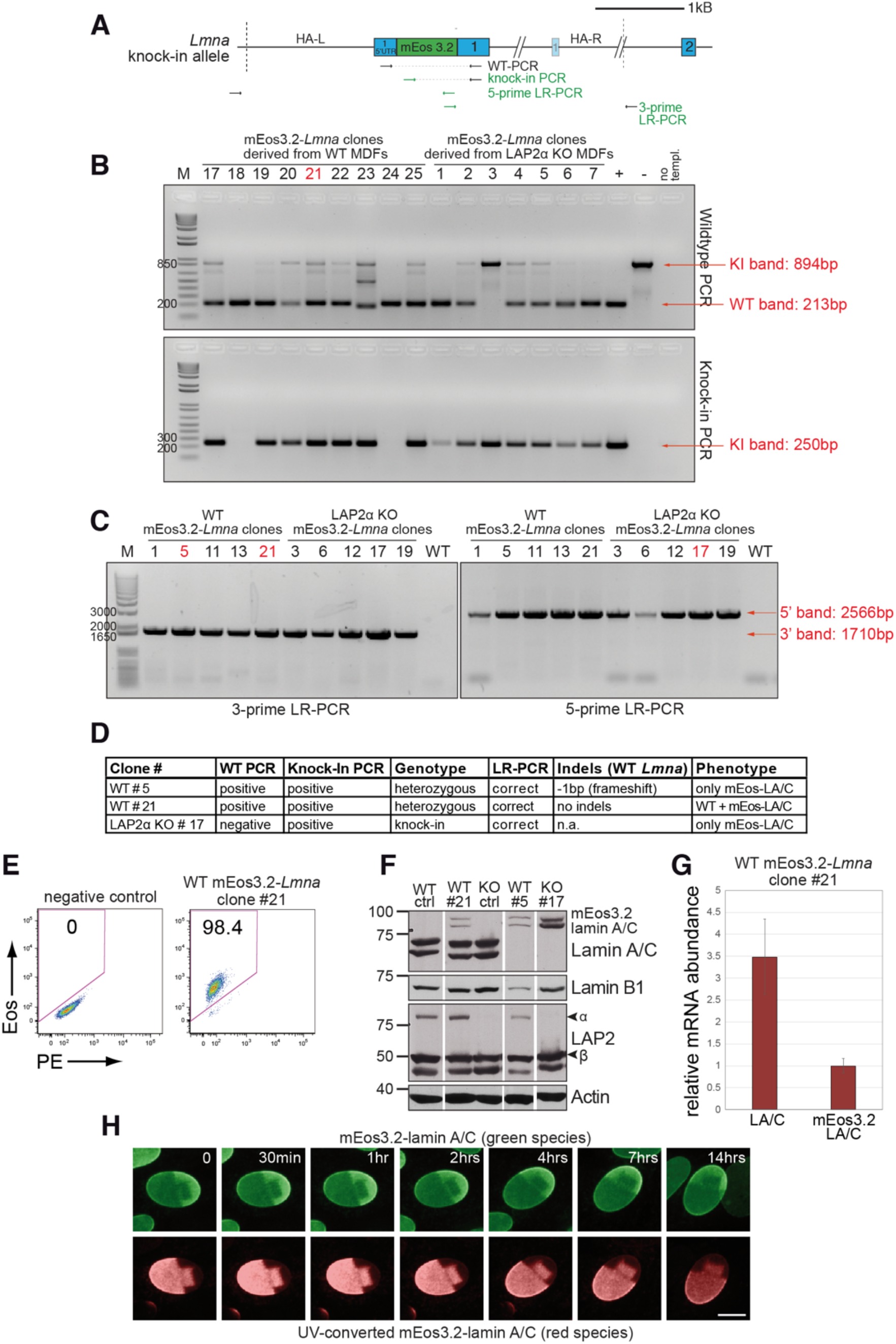
Characterization of mEos3.2-*Lmna* mouse fibroblasts. **(A)** Schematic view of exons 1 and 2 (bars) and adjacent introns (lines) of the mouse *Lmna* locus after integration of mEos3.2. Very long introns are not displayed in their original length as indicated by a double slash. The second, light-colored exon 1 encodes the N-terminus of meiosis-specific lamin C2. Primers used for genotyping and long-range (LR) PCR are indicated (green: specific for modified allele). Dashed lines demarcate the left and right homology arm used for targeting (HA-L: Homology Arm-Left; HA-R: Homology Arm-Right). **(B)** Genomic DNA was isolated from clones derived from either wildtype (WT) or LAP2α knockout (KO) mouse dermal fibroblasts (MDFs). Samples were processed for PCR using the primers indicated in (A) to determine the clonal genotype. **(C)** Clones, where genotyping indicated the presence of a modified allele, were analyzed by long range (LR-)PCR to test for proper integration of the targeting construct. Primer pairs for the left and right homology arm are indicated in (A), with one primer of each pair outside the homology arms. **(D)** Table summarizes results for relevant clones of PCRs in (B), as well as Indels in the WT allele (if present) as determined by PCR across the sgRNA target sequence, followed by sequencing and analysis using the TIDE software. Clonal phenotypes were determined by Western blotting using antibodies to the indicated antigens (anti lamin A/C 3A6), showing expression of mEos3.2-tagged lamins A and C in the presence (WT#21) or absence (WT#5) of untagged wildtype lamin A/C **(F)**. **(E)** Positive clones were also analyzed for the presence of mEos3.2 by FACS. Representative FACS blots of normal wildtype mouse fibroblasts (negative control) and Eos3.2-*Lmna* WT clone #21 are shown. **(G)** Levels of endogenous untagged and mEos3.2-tagged lamins A and C in RNA isolated from WT clone #21 were determined by quantitative qRT-PCR. Graph displays relative mRNA abundance for each mRNA species, revealing a ratio of approx. 1:3. **(H)** WT#21 cells were exposed to UV light (405 nm) to convert mEos3.2-lamin A/C from the green into the red species. Cells were then tracked for 14 hours by live-cell imaging to test for stable integration of mEos3.2 lamin A/C into the peripheral lamina. No significant mobility of converted lamins in the lamina was observed over the entire imaging time. Pictures display merged Z-stacks to visualize most of the peripheral lamina. Bar: 10 μm.

**Figure S4.**
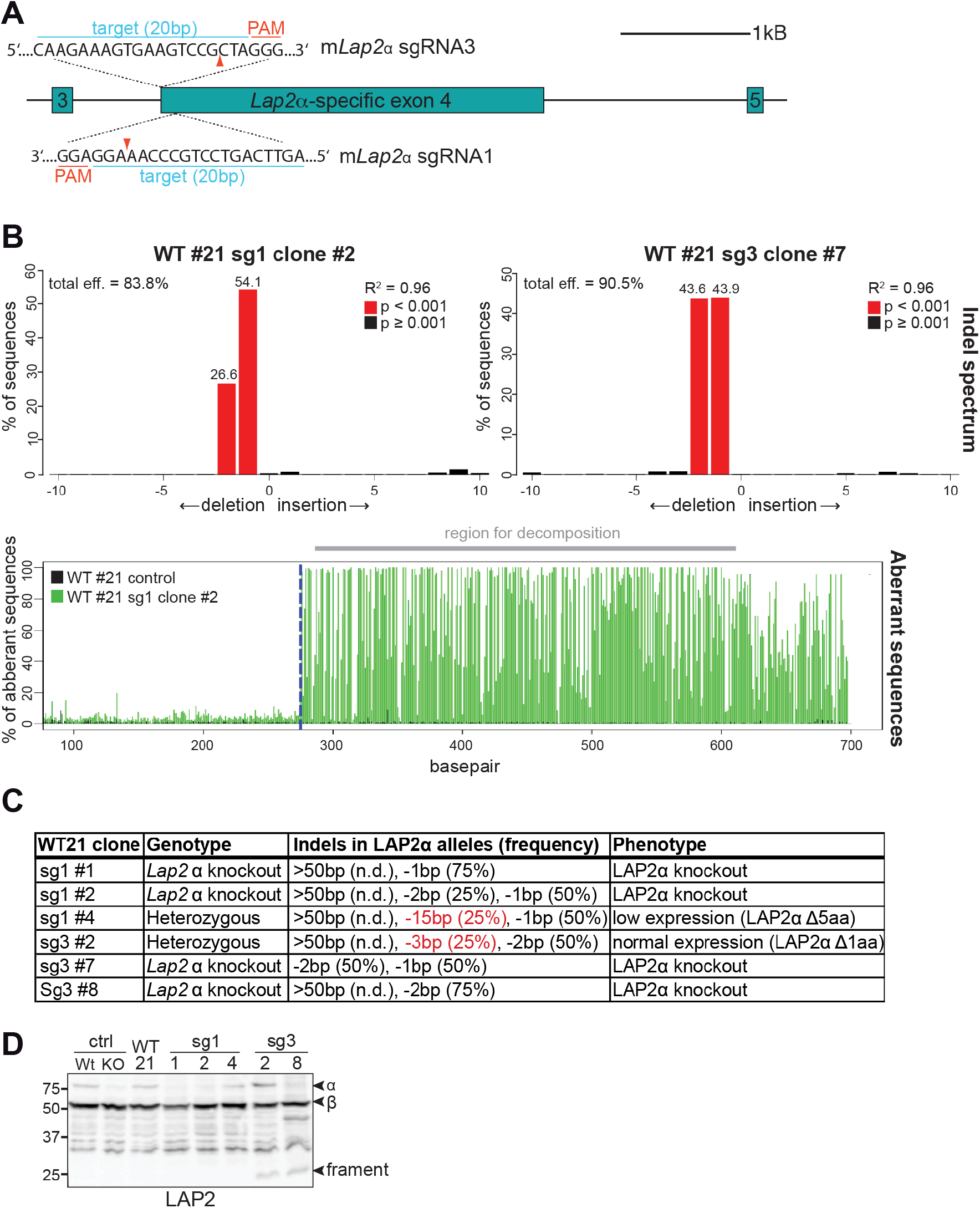
Generation of isogenic LAP2α knockout mEos3.2-*Lmna* clones using CRISPR-Cas9. **(A)** Large interclonal variability in the expression levels of mEos-tagged and untagged lamin A/C in mEos3.2-*Lmna* mouse fibroblasts (Fig. S3) prompted us to use one wildtype clone (WT clone #21) with normal nuclear morphology that harbors both, tagged and untagged lamins A/C, to create isogenic LAP2α knockout cells with two different LAP2α-specific sgRNAs. Image shows a schematic view of exons 3-5 (bars) and adjacent introns (lines) of the murine *Lap2* locus. Positions of the target sequences of *Lap2*α-specific sgRNAs 1 and 3 at the beginning of exon 4 are shown (blue). Protospacer-adjacent motif (PAM) sequences for each sgRNA are marked in red. Red arrowhead; expected Cas9 cut site. **(B)** A region spanning both cut sites was amplified from genomic DNA of the indicated clones and wildtype control cells by PCR. PCR products were sequenced and analyzed using TIDE as described in Fig. S1B. Upper graphs display detected Indels and their frequency in % (numbers atop red bars), lower graph shows an example alignment of wildtype and clone sg1 #2 sequences. Dashed blue line; expected Cas9 cut site. The region used for decomposition of individual sequences and Indel prediction is marked in grey. WT #21; mEos3.2-*Lmna* LAP2α wildtype clone #21 used to generate isogenic knockout lines. **(C)** Summary of generated isogenic mEos3.2-*Lmna* clones and their detected Indels as determined by TIDE analysis. Alleles with deletions >50 bp cannot be analyzed by TIDE. Sequencing revealed that WT#21 and all isogenic clones are tetraploid. Alleles marked in red represent in-frame deletions, encoding full length LAP2α with small deletions. Clonal phenotypes are described based on data in (D). **(D)** Isogenic clones were processed for Western blot analysis using antibodies against the LAP2 common N-terminal domain. Notably, clones generated with LAP2α-specific sgRNA3 express a small N-terminal fragment likely representing the LAP2 N-terminal domain common to all LAP2 isoforms. To control for any effects caused by this LAP2 fragment, sg3 LAP2α knockout clones were not only compared to their wildtype origin clone (WT#21), but also to a clone (sg3 #2) that carried one *Lap2*α allele with an in-frame deletion of 3bp, thus expressing full length LAP2α (minus 1 amino acid), as well as the N-terminal fragment from the other, modified alleles, thus serving as an additional “wildtype” control.

**Figure S5.**
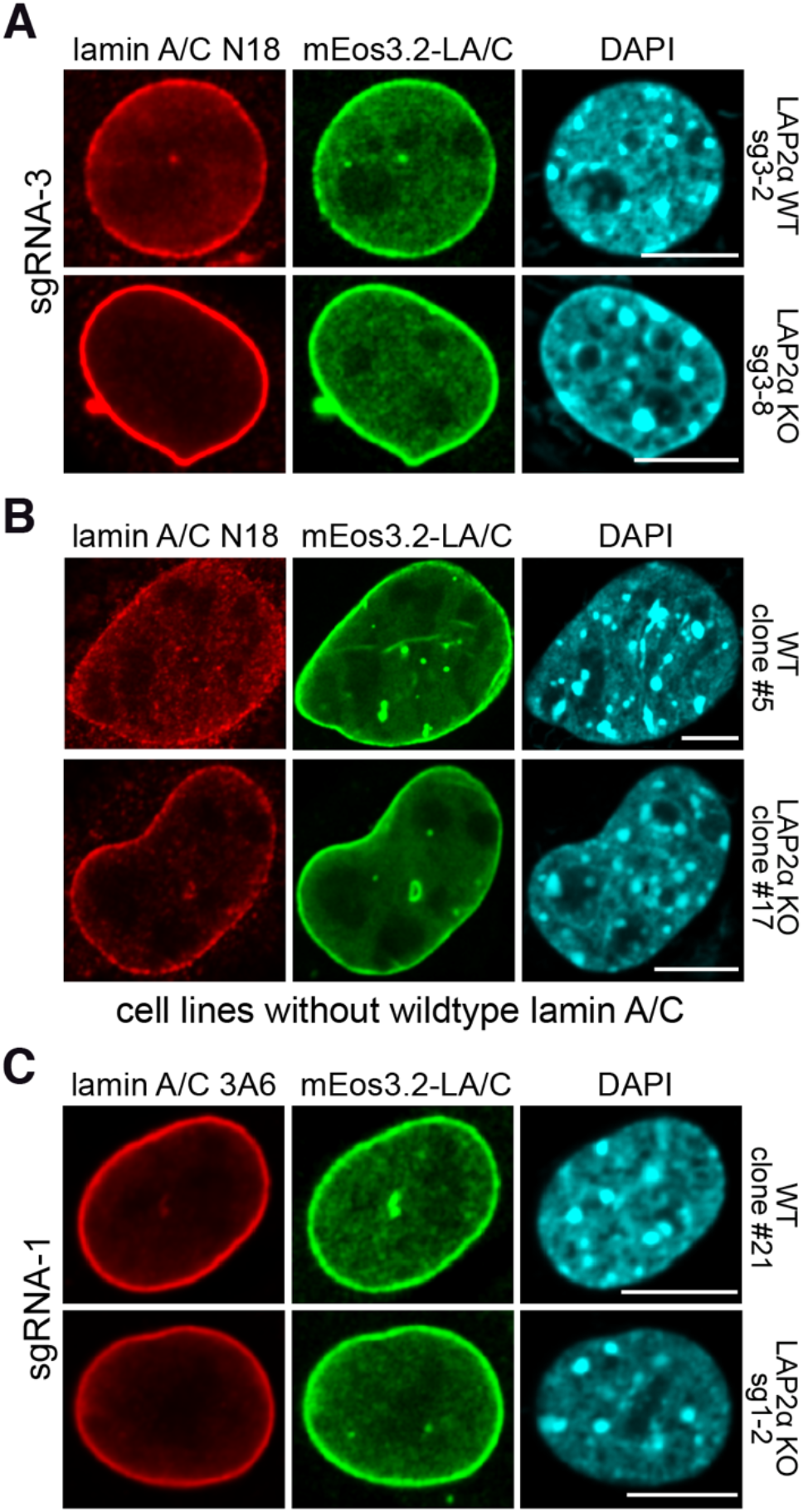
Nucleoplasmic lamin A/C antibody staining but not the mEos3.2 fluorescence signal is reduced in LAP2α knockout versus wildtype fibroblasts in fluorescence microscopy. **(A)** mEos3.2-*Lmna* WT clone sg3-2 and LAP2α KO sg3-8 cells were processed for immunofluorescence microscopy using lamin A/C antibody N18 staining nucleoplasmic lamin A/C, and DAPI to visualize DNA. **(B)** mEos3.2-*Lmna* WT clone #5 and LAP2α KO clone #17 expressing only mEos3.2-tagged lamin A/C without untagged, wildtype lamin A/C (see Fig. S3F), were processed as in (A). **(C)** mEos3.2-*Lmna* WT clone #21 and LAP2α KO sg1-2 cells were processed as in (A) using lamin A/C antibody 3A6 staining mostly peripheral lamin A/C in both LAP2α wildtype and knockout cells. Confocal images are shown in (A), (B) and (C). Bar: 10 μm.

**Figure S6.**
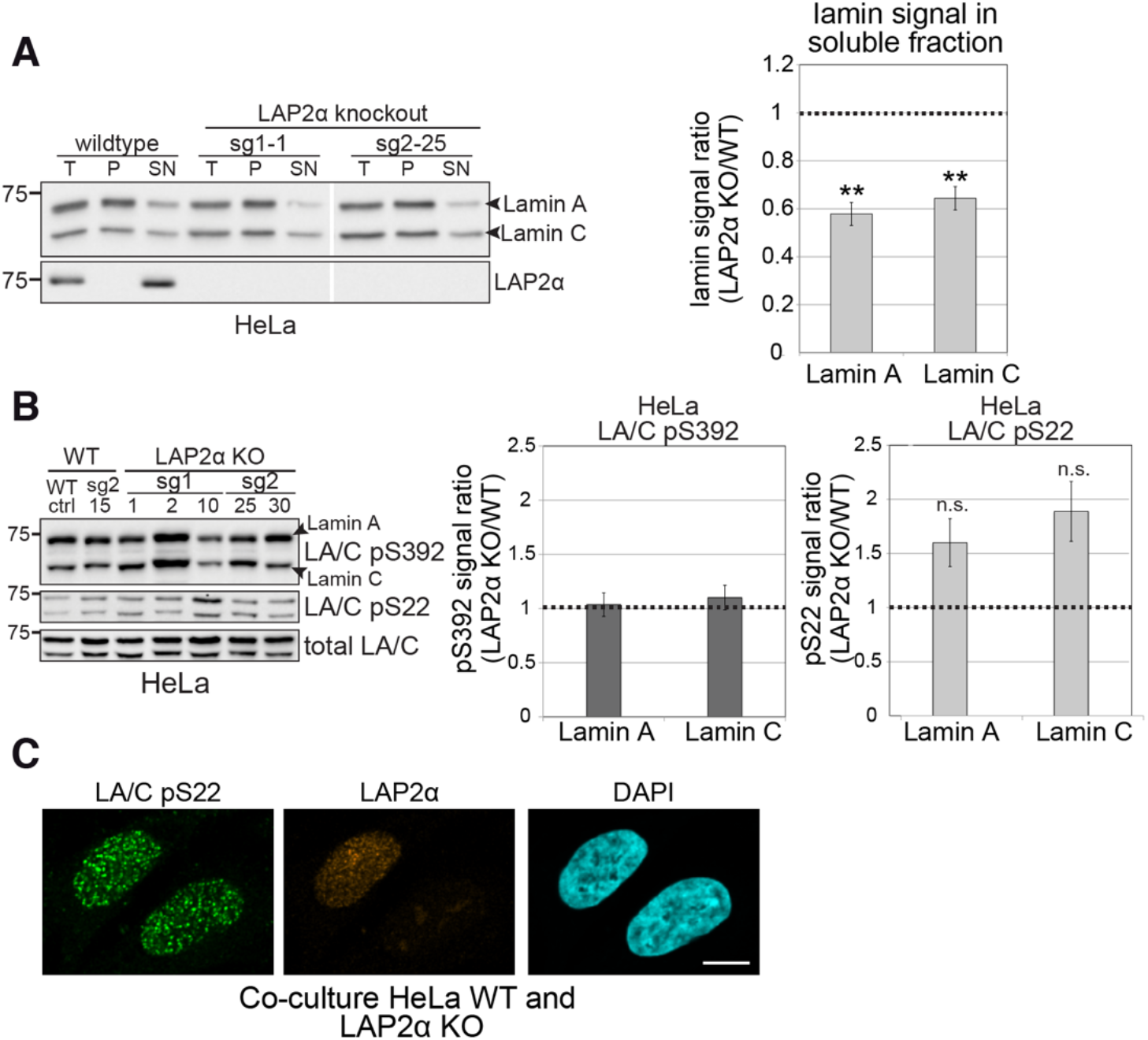
Nucleoplasmic lamins A and C are more resistant to extraction in the absence of LAP2α in HeLa cells without detectable changes in lamin phosphorylation. **(A)** Wildtype and LAP2α knockout HeLa cells were extracted in salt and detergent-containing buffer (150 mM NaCl, 0.5% NP-40). Extracts were processed for Western blot analysis. Left panel shows a representative Western blot of wildtype cells and two LAP2α knockout clones using antibodies against lamin A/C (E1) or LAP2α (Ab15-2). T: total lysate; P: insoluble pellet fraction; SN: soluble, extracted supernatant fraction. Western blots were quantified and lamin signal in the supernatant was normalized to total lamin A/C signal. Graph displays lamins A and C levels in the supernatant fraction of LAP2α knockout samples as average fold difference ± S.E.M over wildtype samples. n_WT_=3, n_KO_=6; **p < 0.01 (p_LA_=0.0073, p_LC_=0.0077; unpaired, two-tailed student’s t test). **(B)** LAP2α KO or WT HeLa clones were processed for Western blotting using antibodies against lamin A/C phosphorylated at specific residues as indicated or a pan-lamin A/C antibody (E1). Western blot signals for phosphorylated lamins were quantified, normalized to total lamins A and C and expressed as fold difference to the WT samples (graphs on the right). Graphs display average fold difference ± S.E.M. n_WT_=2, n_KO_=5, n.s.: non-significant (p_LA_=0.15, p_LC_=0.11; unpaired, two-tailed student’s t test). **(C)** LAP2α KO or WT HeLa clones were co-cultured and processed for immunofluorescence microscopy using antibodies specific to lamin A/C phosphorylated at serine 22 (LA/C pS22) or LAP2α (1H11), and DAPI to visualize DNA. Bar: 10 μm.

**Figure S7.**
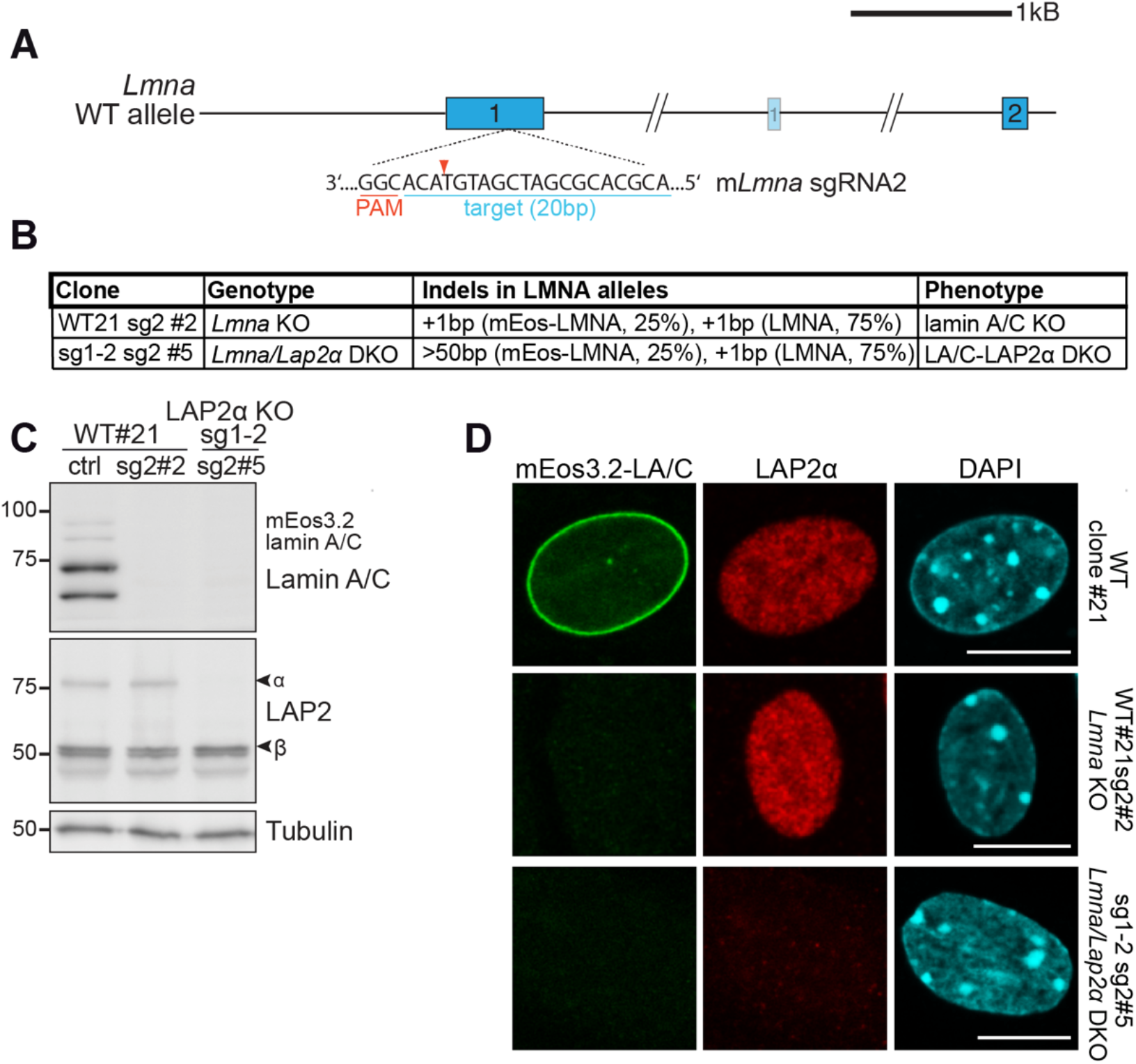
Generation of isogenic*Lmna* knockout and *Lmna/Lap2*α double knockout mouse fibroblasts using CRISPR-Cas9. **(A)** Schematic view of exons 1 and 2 (bars) and adjacent introns of the mouse *Lmna* locus. Very long introns are not displayed in their original length as indicated by a double slash. The second, light-colored exon 1 encodes the N-terminus of meiosis-specific lamin C2. The target sequence of *Lmna*-specific sgRNA2 in exon 1 is shown (blue). Protospacer-adjacent motif (PAM) is marked in red. Red arrowhead; expected Cas9 cut site. **(B)** Summary of generated isogenic *Lmna* knockout clones (using mEos3.2-*Lmna* WT#21 cells) and *Lmna/Lap2*α double knockout clones (using mEos3.2-*Lmna* LAP2α knockout sg1-2 cells) and their detected Indels as determined by TIDE analysis. Alleles with deletions >50 bp cannot be analyzed by TIDE. Clonal phenotypes are described based on data in (C). **(C)** Isogenic clones were processed for Western blot analysis using antibodies to lamin A/C (E1), the LAP2 common N-terminal domain and γ-tubulin as loading control. **(D)** Isogenic clones and mEos3.2-*Lmna* WT#21 control cells were processed for immunofluorescence microscopy using antibodies against LAP2α (1H11) and DAPI to stain DNA and analyzed using an LSM 710 confocal microscope. Bar: 10 μm.

## Video legends

**Video 1.** Wildtype HeLa cells were transiently transfected with EGFP-pre-lamin A and analyzed by live-cell imaging 5 hours post-transfection. Shown is a video assembled from images obtained every 20 minutes of cells starting to display emerging EGFP-pre-lamin A fluorescence (associated with Fig. 1A).

**Video 2.** LAP2α knockout HeLa cells (clone sg1#2) were transiently transfected with EGFP-pre-lamin A and analyzed by live-cell imaging 5 hours post-transfection. Shown is a video assembled from images as described for Video 1 (associated with Fig. 1A).

**Video 3.** Wildtype HeLa cells were transiently transfected with EGFP-pre-lamin A and analyzed by live-cell imaging 24 hours post-transfection. Shown is a video assembled from images obtained every 20 minutes of cells expressing matured EGFP-pre-lamin A starting from a cell undergoing mitosis (associated with Fig. 1B).

**Video 4.** LAP2α knockout HeLa cells (clone sg1#2) were transiently transfected with EGFP-pre-lamin A and analyzed by live-cell imaging 24 hours post-transfection. Shown is a video assembled from images as described for Video 4 (associated with Fig. 1B).

**Video 5.** Wildtype HeLa cells were transiently transfected with EGFP-mature lamin A and analyzed by live-cell imaging 5 hours post-transfection. Shown is a video assembled from images obtained every 20 minutes of cells starting to display emerging EGFP-mature lamin A fluorescence (associated with Fig. S2A).

**Video 6.** Wildtype HeLa cells were transiently transfected with EGFP-ΔK32 pre-lamin A and analyzed by live-cell imaging 5 hours post-transfection. Shown is a video assembled from images obtained every 20 minutes of cells starting to display emerging EGFP-ΔK32 pre-lamin A fluorescence (associated with Fig. S2A).

**Video 7.** Wildtype HeLa cells were transiently transfected with EGFP-ΔK32 mature lamin A and analyzed by live-cell imaging 5 hours post-transfection. Shown is a video assembled from images obtained every 20 minutes of cells starting to display emerging EGFP-ΔK32 mature lamin A fluorescence (associated with Fig. S2A).

**Video 8.** Wildtype HeLa cells were transiently transfected with EGFP-mature lamin A and analyzed by live-cell imaging 24 hours post-transfection. Shown is a video assembled from images obtained every 20 minutes of cells expressing mature EGFP-lamin A starting from a cell undergoing mitosis (associated with Fig. S2B).

**Video 9.** Wildtype HeLa cells were transiently transfected with EGFP-ΔK32 pre-lamin A and analyzed by live-cell imaging 24 hours post-transfection. Shown is a video assembled from images obtained every 20 minutes of cells expressing matured EGFP-ΔK32 pre-lamin A starting from a cell undergoing mitosis (associated with Fig. S2B).

**Video 10.** Wildtype HeLa cells were transiently transfected with EGFP-ΔK32 mature lamin A and analyzed by live-cell imaging 24 hours post-transfection. Shown is a video assembled from images obtained every 20 minutes of cells expressing mature EGFP-ΔK32 lamin A starting from a cell undergoing mitosis (associated with Fig. S2B).

## Notes

### Competing Interest Statement

The authors have declared no competing interest.

